# Multi-level monitoring of EME1-MUS81 in CPT induced nucleolar rDNA repair

**DOI:** 10.1101/2022.04.11.487769

**Authors:** Karishma Bakshi, Easa Nagamalleswari, Jan Breucker, Nathalie Eisenhardt, Richa Tiwari, Román González-Prieto, Chronis Fatouros, Viduth K Chaugule, Heekyoung Lee, Dragana Ilic, Fredrik Trulsson, Gerhard Mittler, Alfred C.O. Vertegaal, Andrea Pichler

## Abstract

We identified EME1, the regulatory subunit of the structure-specific endonuclease complex EME1-MUS81, as substrate for the sumoylated UBC9 and demonstrated synergistic functions in promoting Camptothecin (CPT)-induced nucleolar ribosomal DNA (rDNA) repair (Nagamalleswari et al, co-submitted). Sumoylation of EME1 appears complex involving mono- and poly-sumoylation. Hence, we addressed here whether these modifications differentially regulate EME1 functions by analyzing EME1-variant expressing cell lines including mono-sumo(1)ylation and poly-sumo(2)ylation mimetic fusions. We complemented our analysis with the regulated endogenous EME1 interactome and our observation that Trichostatin A (TSA) induced EME1 and UBC9 sumoylation. Our findings are substantiated by identifying several regulatory proteins and by detecting CPT-induced endogenous di- and poly-sumoylated EME1 in different cell fractions. Together, our data suggest that Histone H4 acetylation, two mono-sumoylation events, poly-sumoylation, ISG15 and ubiquitination sequentially and in mutual dependence tightly control the intracellular localization, recruitment to DNA lesions, enzymatic activity, withdrawal from DNA lesions and the stability of EME1.

## Main

The essential meiotic structure-specific endonuclease 1 (EME1) is the regulatory subunit of the structure-specific endonuclease complex EME1-MUS81, which functions in recombination repair by cleaving nicked Holliday junctions (HJ) and replication forks (RF)^1–3^. We observed EME1 as a substrate of the sumoylated form of UBC9 (S*UBC9) and uncovered synergistic survival functions in Camptothecin (CPT)-induced homology repair in rDNA (Nagamalleswari et al, co-submitted). rDNA is highly repetitive, rich in secondary structures, and highly transcribed, making it one of the most fragile genomic loci in the cell and thus functions as cellular stress sensor. rDNA is located in the nucleolar organizer regions (NOR) of the five acrocentric chromosomes around which nucleoli are formed^4, 5^.

CPT is a natural drug that interferes with the activity of Topoisomerase 1 (Top1), leaving Top1 covalently bound on DNA (Top1cc) adjacent to a nick in the DNA^6–8^. Top1ccs constitute obstacles to the transcription and replication machineries, resulting in collisions between transcription and replication, while nicked DNA upon replication fork “run-off” results in single-ended double strand breaks (seDSBs). These particular breaks cannot be repaired by non-homologous end joining (NHEJ) due to the lack of the second DNA end and therefore rely on homologous recombination (HR) for repair^9, 10^.

Following CPT treatment, the EME1-MUS81 complex induces double-strand breaks (DSBs) as a prerequisite for replication restart^11^, and concordantly both subunits were among the top hits counteracting CPT-induced cell damage^12^. This is of great interest because CPT-based drugs are used in clinical anti-cancer therapy to treat various solid tumors but have the disadvantage of acquired or de novo clinical resistance. Understanding the molecular mechanism leading to this resistance will open up new avenues to improve tumor therapy^6–8^.

We found that CPT exposure and sumoylation promote the accumulation of EME1 and UBC9 in nucleolar repair condensates, suggesting a cooperative role in rDNA repair (Nagamalleswari et al, co-submitted). Homology-repair in repetitive rDNA is error prone, as it can lead to loss of repeats or chromosomal rearrangements that adversely affect cell viability^13^. We proposed a model in which EME1-MUS81 induces DSBs after resection and first template switching for D-loop formation (Nagamalleswari et al, co-submitted). In the case of seDSBs, the resulting structures resemble nicked HJ, the natural substrates of EME1-MUS81^1, 14, 15^. Resolution at this stage prevents the risky second template switch in repetitive rDNA and enables replication through the converging replication fork, analog to MUS81 dependent repair of repetitive Alu elements upon replication fork breakage^16^. Consistent, MUS81 activity is required for maintaining the number of rDNA repeats through replication fork quality control^17^ and sumoylation of EME1 and UBC9 enhanced cell viability (Nagamalleswari et al, co-submitted).

SUMO can regulate protein functions by covalent (sumoylation) attachment or by non-covalent interactions via so-called SUMO interaction motifs (SIMs). Sumoylation depends in most cases on the hierarchical action of E1, E2 and E3 enzymes, but exceptions allow modification with only the E1 and E2^18^. We identified that sumoylation of the single SUMO E2 UBC9 can enhance modification of selected SUMO substrates with a SIM in close distance to the SUMO modification site^19^. SUMO exists in several flavors, of which the ubiquitously expressed SUMO1 and the almost identical SUMO2/3 are the most prominent paralogs (Pichler et al., 2017). A substrate can be modified with a single SUMO (mono-sumoylation), with multiple SUMOs (multi-sumoylation) or with a SUMO chain (poly-sumoylation) preferentially assembled by SUMO2/3s. Our initial analysis revealed that EME1 sumoylation is complex and might involve mono- and multi-/poly-sumoylation as well as SUMO1 and SUMO2/3 (Nagamalleswari et al, co-submitted).

In this study, we aimed to investigate whether mono- and poly-sumoylation differentially regulate EME1 function. To address this, we examined mono-sumo(1)ylation and poly-sumo(2)ylation mimetic, linear EME1 fusion expressing cell lines alongside to additional EME1 variants. We aligned key findings with the endogenous protein, complemented our analysis with the regulated EME1 interactome and our observation that trichostatin A (TSA) enhances both UBC9 and EME1 sumoylation. Our analysis showed that EME1 is tightly regulated at multiple levels, and we uncovered several of its regulators. Finally, we showed that endogenous EME1 is di-and poly-/multi-sumoylated in different cellular compartments. The insights gained allow us to design a model of how EME1 is sequentially and interdependently recruited to sites of DNA damage to introduce DSBs and restart replication before being removed. These molecular insights lay the ground for novel possibilities to inhibit EME1 function to improve CPT-based cancer therapy.

## Results

### EME1 mono- and poly-sumoylation promote cell survival upon CPT exposure

We identified EME1 as largely mono-sumoylated substrate for the sumoylated form of UBC9 and mapped Lysine 27 as major site *in vitro* (Nagamalleswari et al, co-submitted). However, the corresponding mutation did not display a biological phenotype, as it was still heavily sumoylated with SUMO1 and SUMO2/3 in cells. Consistent with this, we confirmed that different SUMO E3 ligases can also modify EME1 *in vitro*, presumably by involving mono-, multi-, and poly-sumoylation (Nagamalleswari et al, co-submitted). In all scenarios tested, EME1 K27 was the major site of modification. Mutation of this site resulted in modification of minor sites, which become more prominent (Nagamalleswari et al, co-submitted). Also in cells, mapping of EME1 SUMO sites is challenging. Conventional mass spec approaches using trypsin digestion resulted in the detection of multiple EME1 sumoylation sites (lysines 159, 232, 241, 340, 357, 441) after heat shock and MG132-induced proteasome inhibition ^20^. Under these conditions, the peptide containing K27 could not be detected because it neutralizes upon sumoylation. Though, in a less sensitive, alternative approach, EME1 K27 was detected as single sumoylation site upon heat shock^21^ confirming it as major site in cells. Taken together, these data suggest that EME1 is likely mono- multi-, and poly-sumoylated in a stress-regulated manner, whereas site specificity appears less important. For UBC9, we demonstrated biochemically that N-terminal sumoylation can be functionally-mimicked by a N-terminal linear SUMO fusion (Nagamalleswari et al, co-submitted). We therefore generated U2OS cell lines that are doxycycline (dox)-inducible and express non-cleavable N-terminal HA-Strep-GFP-tagged (^T^) ^T^EME1-variants: Wild-type (^T^E), ^T^SUMO1-EME1 fusion mimicking mono-sumoylation (^T^S1E) and fusion of ^t^tetra-SUMO2-EME1 mimicking modification with a SUMO2 chain (^T^4S2E). The different ^T^EME1-variants are graphically depicted in Fig. 1a. Because we observed cooperative functions of EME1 and sumoylated UBC9 in CPT response, we analyzed the following treatments throughout the manuscript after induced expression of the respective ^t^EME1-variant: mock, CPT (10 µM) for one hour, and pre-treatment with the SUMO E1-inhibitor ML792 (1 µM) one hour before CPT exposure (CPT & ML792). These treatments allows us to distinguish between sumoylation and non-sumoylation-dependent CPT-induced EME1 functions. In selected experiments, we also performed ML792 treatment (1 µM for 2 hours) alone. Co-Immunoprecipitation analysis, shown in Fig. 1b (Input in Extended Data Fig. 1a), indicates that ^T^E and ^T^4S2E are expressed at much lower levels than ^T^S1E.

**Fig. 1.**
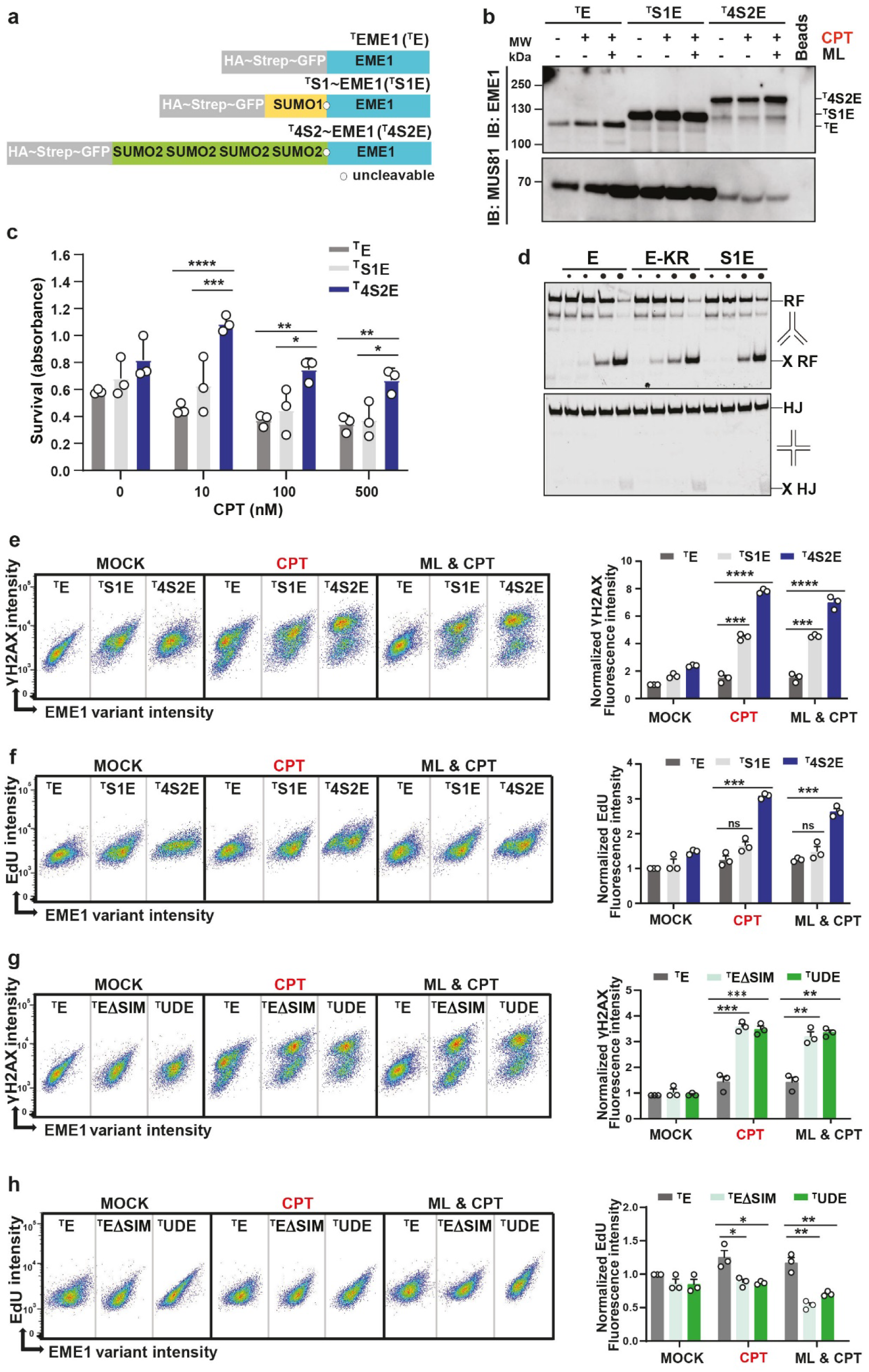
EME1 mono- and poly-sumoylation promote cell survival, DSBs & replication upon CPT exposure. **(a)** Schematic representation of HA-Strep-GFP tagged full-length EME1-variants: ^T^E wild-type, mono-sumoylated SUMO1 fusion ^T^S1E, and poly-sumoylated SUMO2 fusion ^T^4S2E, that were stable integrated in the FRT locus of U2OS cells for doxycycline (Dox) inducible expression. ° uncleavable. **(b)** Immunoblots of GFP-Trap enriched ^T^E-variants and their interactions with MUS81. ^T^E, ^T^S1E, and ^T^4S2E expression were induced for 16 hours before treatments with DMSO (vehicle control, MOCK), ML (ML792; 1 µM for 2 hours), CPT (10 µM for 1 hour) or ML & CPT. GFP-Trap-enriched samples were separated on SDS-PAGE and detected with anti-EME1(serum B) or anti-MUS81 antibodies. Control was with GFP-Trap beads. Input control is shown in Extended Data Fig. 1a. **(c)** Diagram of clonogenic survival assay. ^T^E, ^T^S1E, and ^T^4S2E expression were induced for 16 hours before treatments with indicated concentrations (0, 10, 100 and 500 nM) of CPT for one hour. Colonies were stained with 0.5% Crystal Violet after 5-7 days. Images were analyzed with ImageJ and absorbance was measured at 594nm. Bars show the average (+SEM) surviving colonies of three different experiments in triplicates. Differences between ^T^E, ^T^S1E, and ^T^4S2E expressing cells are significant based on two-way ANOVA by Graphpad Prism analysis (*: p-value < 0.05; **: p-value < 0.001, ***: p-value < 0.0005, ****: p-value < 0.0001). Bar chart normalized to ^T^EME1 variant’s mock is shown in Extended Data Fig. 1b (**d)** Endonuclease assay with different DNA substrates. DNA, DY782-end-labeled on one oligonucleotide, was incubated with increasing concentrations of indicated MUS81-EME1 variants (190, 380, 760, 1520 pM) (Upper panel). RF cleavage was monitored after 60 min at 37 °C. Samples were analyzed by native PAGE. RF: replication fork; X RF: cleaved replication fork; EME1 (E); SUMO1-EME1 (S1E); EME1-K27/28R (E-KR). As in Upper panel, but with X12 Holliday junction as DNA substrates (HJ). HJ: Holliday junction; X HJ: cleaved Holliday junction (Lower panel). **(e)** Quantitative analysis of DNA damage upon CPT exposure. Asynchronously grown population of U2OS cells induced for expression of the indicated ^T^EME1 variant (^T^E, ^T^S1E and ^T^4S2E) before treatment with DMSO (MOCK), CPT (10 µM for 1 hour) or ML & CPT (ML792 1 µM for 2 hours) in the presence of 100 µM EdU for 1 hour, fixed and stained with DAPI, γH2AX, and EdU. Scatter plots (left panel) depict cell populations with each dot presenting a single cell. γH2AX fluorescent intensities are shown as function of ^T^EME1-variant (^T^E, ^T^S1E and ^T^4S2E) fluorescent intensities. Bar charts (right panel) show average values of γH2AX levels of three different experiments in triplicates for the indicated cell lines and conditions. Values were normalized using mock of ^T^E conditions. Differences between ^T^E, ^T^S1E and ^T^4S2E expressing cells are significant based on two-way ANOVA followed by Tukey test for multiple comparisons Graphpad Prism 8.0 analysis (*: p-value < 0.05; **: p-value < 0.001, ***: p-value < 0.0005, ****: p-value < 0.0001). **(f)** Quantitative analysis of replication. As **(e)** but scatter plots depict EdU signals as function of ^T^EME1 variant fluorescent intensities. Bar charts (right panel) show averages of EdU levels of three different experiments in triplicates for the indicated cell lines and conditions. **(g)** Quantitative analysis of DNA damage as in **(e)** but with ^T^EΔSIM and ^T^UDE expressing cells. Schematic representation and immunoblots of dox induced expression are shown in Extended Data Fig.1c for ^T^UDE and in Extended Data Fig1d for ^T^EΔSIM. **(h)** Quantitative analysis of replication as in **(f)** but with ^T^EΔSIM and ^T^UDE expressing cells.

Importantly, all EME1 variants interact with MUS81 depending on their abundance and are thus able to form active endonuclease complexes. However, ^T^4S2E-MUS81 interaction was reduced. When we turned to the treatments, mainly ^T^E appeared to be regulated, slightly stabilized upon CPT exposure and further enriched upon ML792 & CPT co-treatments. To sum up, all ^T^EME1-variants can form endonuclease complexes with MUS81. Mono-sumoylation appears to have stabilizing functions.

Transient replacement studies have shown that S1E provides a survival advantage upon CPT exposure (Nagamalleswari, et al. co-submitted). We now aimed to verify whether this was also the case for the stable cell lines and how poly-sumoylation compared to them. As can be seen in Fig. 1c and Extended Data Fig. 1b, both sumoylation variants provided a survival advantage under all conditions tested, with poly-sumoylation having greater effects than mono-sumoylation, even in mock treated cells. This raised the question, whether sumoylation promotes the nuclease activity of the EME1-MUS81 complex, which is beneficial for cell survival upon CPT induced DNA damage. To test this, we compared recombinant EME1-MUS81 (E), EME1^K27/28R^-MUS81 (E-KR) and S1^∼^EME1-MUS81 (S1E) complexes for their abilities in Replication forks (RF) cleavage (Fig. 1d, upper panel) and Holliday junction (HJ) resolution (Fig. 1d, lower panel). However, we did not detect significant differences between all tested complex variants, suggesting that sumoylation does not regulate the catalytic endonuclease activity per se.

### EME1 mono- and poly-sumoylation induce DSBs and poly-sumoylation promotes replication

Earlier studies have shown that EME1-MUS81 generates DSBs upon CPT exposure, which serve as a prerequisite for replication restart and cell survival^11^. Thus, we examined ^T^EME1 variant-expressing cells for their ability to induce DSBs and replication upon mock, CPT and ML792 & CPT treatments. We performed Fluorescence-Activated Cell Sorting (FACS) analysis by co-staining the cells for GFP (^t^EME1 variant expression), γH2AX (Serine139-phosphorylated histone H2AX) as marker for DSBs, EdU (5-ethynyl-2’-deoxyuridine) incorporation to monitor replication, and DAPI for the DNA content. We analyzed γH2AX (Fig. 1e) and EdU (Fig. 1f) as a function of ^T^EME1 variant expression, respectively. Scatter plots are shown on the left and bar charts with average signals of the indicated cell lines and treatments normalized to ^T^E values are displayed on the right. Our analysis from three experiments in triplicates indicated that ^T^S1E significantly improved DSB formation upon CPT exposure, largely independent on additional sumoylation events (CPT & ML792). ^T^4S2E reinforced this effect further but this was to some extent depending on additional sumoylation. When we turned to replication, we observed statistically significant enhancement only with ^T^4S2E, again partially dependent on additional sumoylation events. Together, these data indicate that mono-sumoylation and, even better, poly-sumoylation promote cell survival, independent of direct targeting the nuclease activity of the EME1-MUS81 complex. However, both modifications increase DSB formation to varying degrees, while poly-sumoylation enhances replication.

These findings would greatly benefit from the availability of a EME1-“loss of sumoylation” cell line. Since we observed that sumoylation jumps upon mutation to adjacent sites, we alternatively fused the catalytic domain of the yeast SUMO isopetidase Ulp1 N-terminally to EME1 to prevent its sumoylation^22^. In addition, we have included an EME1 variant mutated in its first SUMO interaction motif (SIM), which is essential for enhanced sumoylation with the sumoylated UBC9 (Nagamalleswari, et al). According to the other EME1-variant lines, we generated a Ulp1-Domain-EME1 expressing line (^T^UDE) and an EME1 line mutated in SIM1 (^T^EΔSIM). Graphical representation and induced expression is shown in Extended Data Fig. 1c,d. Subsequent, the same FACS analysis as for the other variants was performed (Fig. 1g, h). Interestingly, both lines showed increased DSB formation concomitant with decreased replication, likely indicative for the accumulation of detrimental DSBs rather than “repair” DSBs.

### Sumoylation signature and SUMO/SIM interaction direct EME1’s subcellular localization

We next examined the intracellular localization of all ^T^EME1-variants by comparing mock, ML792, CPT and ML792 & CPT treatments in immunofluorescence analysis (IFA) using confocal microscopy (Fig. 2 and Extended Data Fig. 2). Representative single-cell images are shown in main figures and corresponding multi-cell images in Extended Data. Surprisingly, in mock and ML792 treated cells, we observed that ^T^E and ^T^4S2E are largely cytoplasmic, while ^T^S1E was nuclear (Fig. 2a-c, first panels and second panels). Though, when we expose cells to CPT, also ^T^E and ^T^4S2E accumulated in the nucleus (Fig. 2a-c, third panels) and in this case it was abrogated by ML792 treatments (Fig. 2a-c, fourth panels). We next asked, whether we can confirm ^T^E localization at endogenous EME1 levels with an antibody raised against full length EME1 in rabbits. As shown in Fig. 2d, this is indeed the case. Like ^T^E, also endogenous EME1 is mainly cytosolic in unstressed conditions but enriched in the nucleus in a CPT and sumoylation dependent manner. Interestingly, when we turned to the EME1 variants ^T^UDE and ^T^EΔSIM, we observed that ^T^UDE was cytosolic (Fig. 2e) and ^T^EΔSIM nuclear (Fig. 2f), in all conditions tested.

**Fig. 2.**
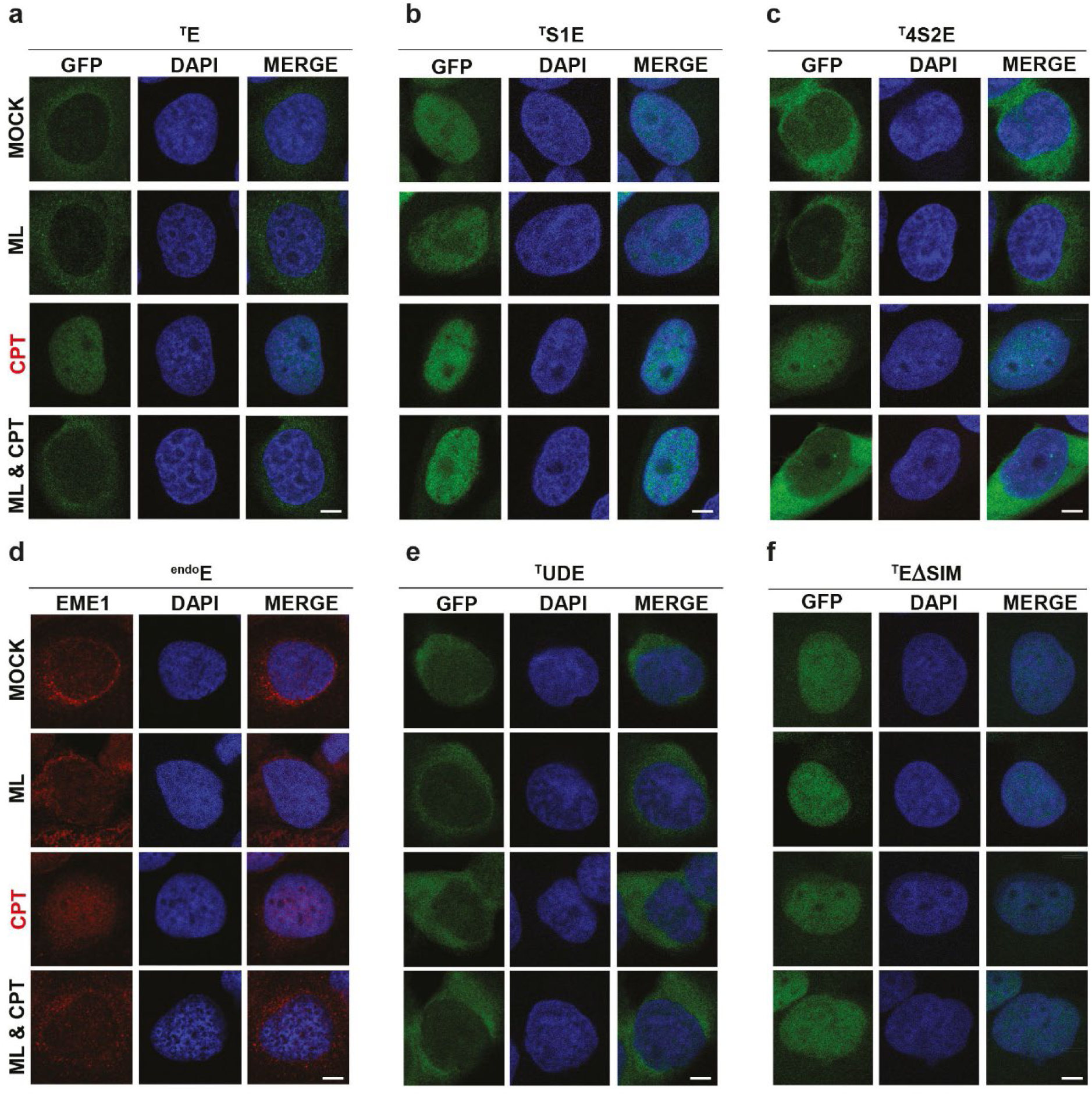
Sumoylation signature and SUMO/SIM interaction direct EME1’s intracellular placements. Immunofluorescence analysis upon induction of ^T^E-variant expression. Treatments with either DMSO (MOCK), ML (ML792; 1 µM for 2 hours), CPT (10 µM for 1 hour) or ML & CPT. Co-staining was with anti-GFP antibody (labelled with Alexa 488, green) for detecting ^T^E-variant and DAPI (blue) for visualization of nuclei. Shown are individual stains and merge of representative single cells. Scale bar represents 10 μm. Related multi cell images are shown in Extended Data Fig. 2a-f. **(a)** Immunofluorescence analysis of ^T^E. **(b)** Immunofluorescence analysis of ^T^S1E. **(c)** Immunofluorescence analysis of ^T^4S2E. **(d)** Immunofluorescence of endogenous EME1 (^endo^E) in U2OS cells but with anti-EME1 serum A. **(e)** Immunofluorescence analysis of ^T^UDE. **(f)** Immunofluorescence analysis of ^T^EΔSIM.

We conclude from this part that EME1 is a shuttling protein. Mono-sumoylation and the lack of the first SIM trapped EME1 in the nucleus, while EME1 wild-type and poly-sumoylated EME1 were largely cytosolic and depend on additional sumoylation events for their nuclear import upon CPT exposure. Loss of sumoylation (^T^UDE) prevented nuclear placement, while the first SIM is likely required for nuclear export.

### Import and export receptors involved in EME1’s intracellular localization

To gain molecular insights into these localization events, we set out to identify the EME1 interactome by performing endogenous EME1 immunoprecipitations (IPs) upon mock, CPT and ML792 & CPT treatments followed by mass spectrometric (MS) analysis. We included IgG IPs as background control. Samples were separated by SDS gel electrophoresis (Fig. 3a), and stained with Coomassie blue. Subsequently each lane was divided into nine fractions (indicated grid), and analyzed individually by MS. Indeed, we found several transport receptors interacting with EME1 in a drug-regulated manner (Fig. 3b).

**Fig. 3.**
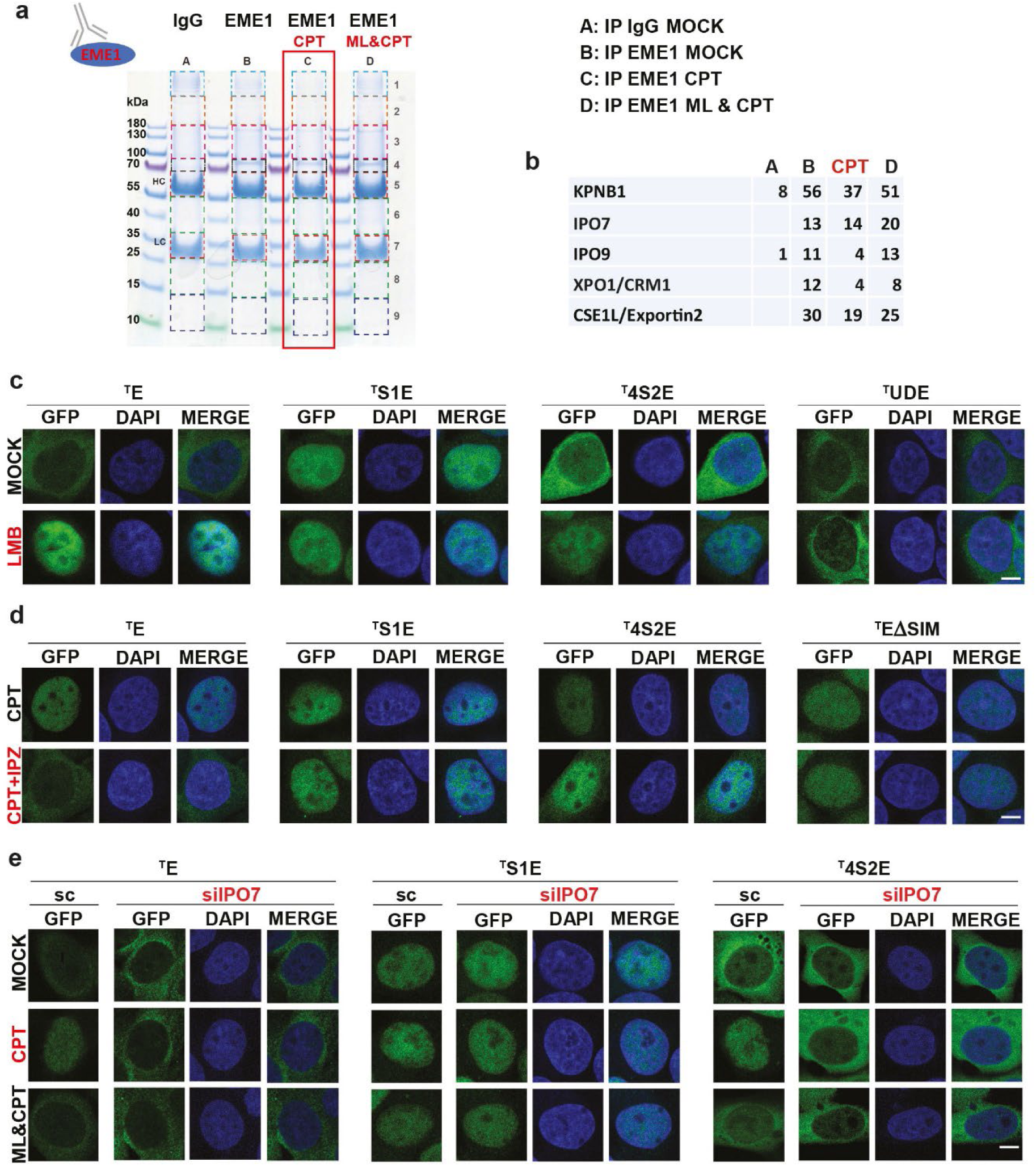
Import and export receptors involved in EME1’s intracellular localization. **(a)** Coomassie blue stained gel of IgG (A) and EME1 Immunoprecipitations (IP) upon mock (B), CPT (C) and ML792 & CPT (D) treatments. Each lane was subdivided into nine fractions (indicated grid) and analyzed individually by mass spectrometry (MS). **(b)** Peptide numbers obtained from EME1 IP-MS for indicated transport receptors upon indicated condition. **(c)** Immunofluorescence analysis of ^T^E-variant expressing (^T^E, ^T^S1E, ^T^4S2E, and ^T^UDE) cell lines upon Leptomycin B (LMB) treatment. Cells were treated with ethanol (MOCK) and 80 nM LMB for 3 hours. Co-staining was with anti-GFP antibody (labelled with Alexa 488, green) for detecting ^T^E-variants and DAPI (blue) for visualization of nuclei. Shown are individual stains and merge of representative single cells. Scale bar represents 10 μm. Related multi cell images are shown in Extended Data Fig. 3a. (**d)** Immunofluorescence analysis as in **(c)** but for ^T^E-variants (^T^E, ^T^S1E, ^T^4S2E, ^T^EΔSIM) upon Importazole (IPZ) treatment. Cells were treated with DMSO (MOCK) and 75 µM IPZ for 2 hours, followed by 1 hour treatment with CPT. Related multi cell images are shown in Extended Data Fig. 3b. **(e)** Immunofluorescence analysis as in **(c)** but for ^T^E-variants (^T^E, ^T^S1E, and ^T^4S2E) upon IPO7 depletion. siRNA mediated knock down with scrambled (sc) or IPO7 (siIPO7) specific siRNAs for 48 hours. Treatments with DMSO (MOCK), CPT (10 µM for 1 hour) or ML (ML792, 1 µM for 2 hours) & CPT. Related multi-cell images for sc are shown in Extended Data Fig. 3c and for siIPO7 in Extended Data Fig.3d, and immunoblots confirming IPO7 knock down in Extended Data Fig. 3e.

As import-receptors, we identified the rather general receptor KPNB1/importin-β^23^, next to IPO7/importin-7 and IPO9/importin-9, receptors implicated in histone and ribosomal protein nuclear import^24, 25^. We found two major export receptors to be CRM1/Exportin-1/XPO1, which exports various cargos^26^, and Exportin-2/CSE1L, the export receptor for importin-α^27^. To confirm these hits, we began our analysis by investigating the established inhibitors Leptomycin B (LMB)^28^ and Importazole (IPZ) that target CRM1 and importin-β functions^29^, respectively. As shown in Fig. 3c, LMB treatment resulted in nuclear localization of ^T^E, ^T^S1E and ^T^4S2E in otherwise untreated cells, while ^T^UDE remained excluded from the nuclei. When we exposed cells to IPZ, which we examined after CPT exposure, we observed that only ^T^E remained in the cytosol, while ^T^S1E, ^T^4S2E and ^T^EΔSIM1 were still able to enter the nucleus (Fig. 3d). Consequently, only ^T^E depends on KPNB1 for nuclear import. Studying IPO7 by siRNA mediated downregulation, we found that both ^T^E and ^T^4S2E required IPO7 for CPT dependent nuclear import (Fig. 3e and Extended Data Fig. 3a-d).

To sum up, we observed that ^T^E and ^T^4S2E are shuttling proteins involving IPO7 for nuclear import and CRM1 for nuclear export. ^T^E requires in addition KPNB1. Under all conditions analyzed, ^T^S1E and ^T^EΔSIM1 remained in the nucleus, though they demonstrated different consequences on CPT induced DSB formation. ^T^UDE was excluded from the nucleus under all conditions tested, suggesting that sumoylation is essential for nuclear import/retention, while SIM1 is required for nuclear export.

### HAT1 and acetylated Histone H4 escort EME1 to DNA lesions

Unexpectedly, when we examined the SUMO2 proteome after treatment with the Histone deacetylase (HDAC) inhibitor Trichostatin A (TSA)^30^, EME1 and UBC9, were amongst the significant hits for increased sumoylation in HeLa cells (Fig. 4a). Hence, we screened the endogenous EME1-interactome (Fig. 3a) for acetylation-associated proteins and identified CPT-promoted interaction with the histone acetyltransferase 1 (HAT1), its substrate histone H4 and all other core Histones (Fig. 4b).

**Fig. 4.**
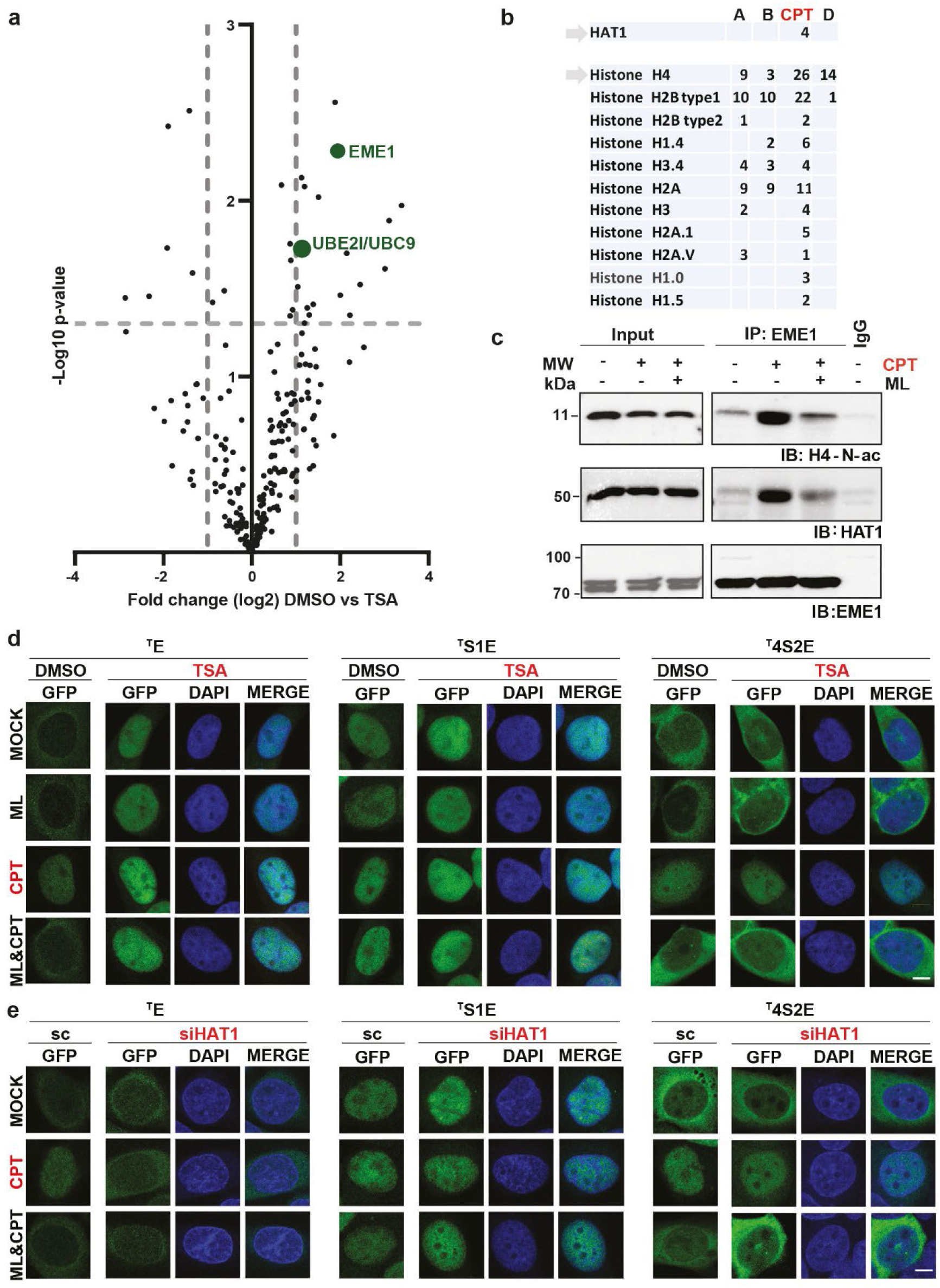
HAT1 and acetylated Histone H4 escort EME1 to DNA lesions. **(a)** Volcano plot of His10-SUMO2 substrates from HeLa cells upon TSA treatment (600nM for 18hrs). Presented are differentially sumoylated proteins amongst His10-SUMO2 expressing cells treated with TSA or DMSO. Dashed lines indicate a cut-off at twofold change (Log2 =1) and a p value of 0.05 (-log10=1.3). EME1 and UBC9/UBE2I are highlighted in green. **(b)** Peptide numbers obtained from EME1 IP-MS analysis (Fig.3a) for HAT1 and various Histones. IgG (A) and EME1 Immunoprecipitations (IP) upon mock (B), CPT (C) and CPT & ML792 (D) treatments. **(c)** Immunoblot monitoring the interaction of EME1 with HAT1 and N-terminal acetylated Histone H4 upon DMSO (MOCK), CPT (10 µM for 1 hour) or ML (ML792, 1 µM for 2 hours) & CPT treatments. Endogenous EME1 (anti-EME1 serum B) or IgG IPs were performed from RIPA buffer extracts. Eluted samples were separated on SDS-PAGE and detection was with anti- H4-N-ac (K5,K8,K12& K16), anti-HAT1 or anti-EME1 (Serum A) antibodies. **(d)** Immunofluorescence analysis of ^T^E-variant (^T^E, ^T^S1E, and^T^4S2E) expressing cells upon Trichostatin A (TSA) treatment. Cells were treated with DMSO (MOCK), or 600 nM TSA for 16 hours, before treatment DMSO, ML, CPT or ML & CPT as in **(c)**. Co-staining was with anti-GFP antibody (labelled with Alexa 488, green) for detecting ^T^E-variants and DAPI (blue) for visualization of nuclei. Shown are individual stains and merge of representative single cells, Scale bar represents 10 μm. Related multi-cell images are shown in Extended Data Fig. 4a. **(e)** Immunofluorescence analysis of ^T^E-variants (^T^E, ^T^S1E, and^T^4S2E) as in **(d)** but upon HAT1 depletion. siRNA mediated knock down with scrambled (sc) or HAT1 (siHAT1) specific siRNAs for 48 hours before treatments with DMSO (MOCK), ML, CPT or ML & CPT as in **(c)**. Related multi cell images are shown in Extended Data Fig.4b and immunoblots confirming HAT1 knock down in Extended Data Fig. 4c.

HAT1 di-acetylates newly synthesized histones H4 on lysines 5 and 12 (H4K5/12ac) in the cytoplasm and escorts their import into the nucleus, where they transiently accompany replication-coupled chromatin maturation. The exact function of H4K5/12ac is unclear, but chromatin maturation begins with deacetylation of H4K5/12ac and HAT1 displacement^31–33^. Upon recovery from replication stress, HAT1 is stabilized at stalled replication forks and protects the newly synthesized DNA from Mre11-mediated degradation^32, 34^. Moreover, HAT1 is required for homology repair but not for NHEJ, and its loss decreases cell viability upon CPT exposure^35^. In *Saccharomyces cerevisiae* Hat1p is recruited to sites of DSBs, where it is responsible for H4 K5 /12 acetylation^36^. Hence, we examined the regulated interaction of EME1 with HAT1 and N-terminal acetylated histone H4 and confirmed the CPT induced binding (Fig. 4c).

We next asked whether TSA exposure or HAT1 depletion was involved in regulating the intracellular localization of EME1-variants. In fact, TSA treatment resulted in nuclear localization of ^T^E (Fig. 4d and Extended Data Fig. 4a) but was in contrast to CPT treatment (Fig. 2a) independent of additional sumoylation events (ML792 & TSA), even in TSA & CPT co-treatments (Fig. 4d, left lower panels). However, ^T^S1E and ^T^4S2E remained unaffected under all conditions tested (Fig. 4d, middle and right panel), suggesting different import routes for modified and unmodified EME1. The same scenario was observed by siRNA-mediated depletion of HAT1 (Fig. 4e and Extended Data Fig. 4b-c). The nuclear localization of ^T^E upon CPT exposure was clearly dependent on HAT1 and sumoylation (Fig. 4e, left panel). Again, both sumoylation-mimetic EME1 variants remained unaffected (Fig. 4e, middle and right panel).

In summary, TSA treatment and HAT1 promotes nuclear import/retention of ^T^E, independent of its sumoylation, while ^T^4S2E and ^T^S1E intracellular localization remains unchanged. We conclude from these results that EME1 is escorted by HAT1 and acetylated H4 histones to DNA and possibly stabilized in the presence of DNA lesions, while its sumoylated versions appear to use other routes to enter the nucleus.

### EME1 is sumoylated and retained at DNA lesions in nucleolar repair condensates

Upon CPT exposure, ^T^E appears to be imported by accompanying HAT1 and H4K5/12ac. However, because its nuclear localization depends on sumoylation, we wondered whether EME1 is stabilized at DNA lesions by sumoylation in the presence of specific DNA structures. To test this possibility, we performed *in vitro* sumoylation assays in the absence and presence of HJ, RFs and single stranded DNA (ssDNA). As shown in Fig. 5a, we indeed observed enhanced EME1 sumoylation exclusively in the presence of the natural EME1 substrates, HJs and RFs, but not with ssDNA. Moreover, only selected enzymes, including sumoylated UBC9^19^, PIAS1 and ZNF451-3^37^, promoted DNA structure-specific sumoylation (Fig. 5a), whereas we did not detect it, neither with the SUMO E3 ligase fragments RanBP2γFG ^38^ nor with ZNF-N^37^. Other DNA structures, such as double stranded DNA (dsDNA) or plasmid DNA (plasmid) also did not result in increased sumoylation. Taken together, these data support the idea that EME1 is sumoylated when bound to DNA in the presence of specific DNA structures and selected SUMO enzymes.

**Fig. 5.**
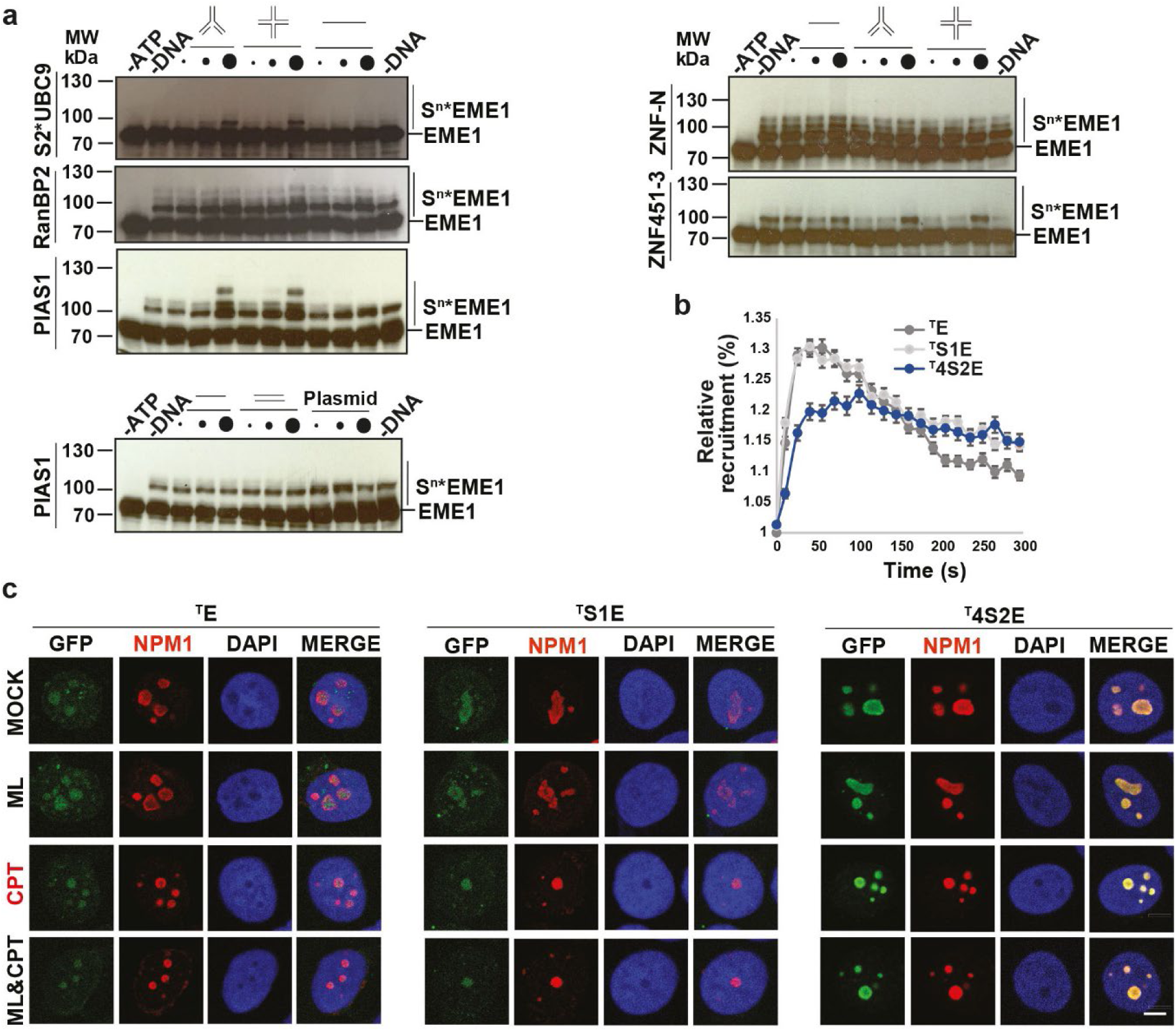
EME1 is sumoylated and retained at DNA lesions in nucleolar repair condensates. **(a)** Immunoblot of DNA dependent *in vitro* sumoylation of EME1. Assay was carried out in the presence of S2*UBC9 (125 nM) or E3 ligases - BP2ΔFG (1 nM), PIAS1 (1 nM), ZNF-N (2 nM) and ZNF451-3 (10 nM) with increasing concentrations (4, 20 and 100 nM) of different forms of DNA - replication fork (RF), Holliday junction (HJ) single stranded DNA (-), double stranded DNA (=), and plasmid and incubated with 200 nM EME1- MUS81 complex, 60 nM E1, 50 nM E2, 2 μM SUMO2 and 5 mM ATP in 20μL reactions for 30 minutes at 30°C. Samples were stopped with SDS sample buffer and analyzed by immunoblotting using anti-EME1 serum A antibodies. **(b)** Recruitment of ^T^E -variants to DNA damage tracks. ^T^E, ^T^S1E, and ^T^4S2E expression was induced, micro-irradiated with a 2-photon laser and recruitment of the tagged protein was followed in a time course. Shown is the relative recruitment quantification of the ^T^E constructs to the laser induced DNA damage tracks. Average and SEMs of three different experiments are shown (N ^T^E =95; N ^T^S1E =107; N^T^4S2E =56). **(c)** Immunofluorescence analysis to monitor ^T^E-variant (^T^E, ^T^S1E, and ^T^4S2E) localization in CSK-buffer insoluble cellular condensates. ^T^E-variant expressing cell lines were treated with DMSO (MOCK), ML (ML792, 1 µM for 2 hours), CPT (10 µM for 1 hour) or ML & CPT, followed by CSK-buffer treatment to release soluble cellular proteins. Staining was with anti-B23 antibody (coupled to Alexa 594, red) for detecting Nucleophosmin (NPM1), and anti-GFP antibody (labelled with Alexa 488, green) for detecting ^T^E-variants. DAPI (blue) was used for visualization of nuclei, and merge shows localization of proteins along with DAPI. Shown are representative single cells. Scale bar represents 10 μm. Related multi cell images are shown in Extended Data Fig. 5a.

To determine whether sumoylation of EME1 alters recruitment or stabilization to DNA lesions, we performed laser microirradiation-induced DNA damage tracks in the EME1-variant lines (Fig. 5b). Recruitment of ^T^E and ^T^S1E to DNA lesion was comparable, whereas ^T^4S2E was delayed. However, over time both, ^T^S1E and ^T^4S2E remained stabilized at DNA lesion, while the signal for ^T^E was already decreasing, indicating its removal.

In our previous work we observed that EME1 accumulates in nucleolar repair condensates after CPT exposure along with UBC9 (Nagamalleswari et al, co-submitted). To this end, we applied a protocol established for the detection of DNA repair condensates by pretreating cells with a mixture of detergent and sucrose (CSK buffer) to release soluble proteins prior to immunofluorescence staining^39, 40^. In these IFA, EME1 was already detectable in nucleolar repair condensates in untreated cells, CPT treatment promoted EME1 enrichment in these nucleolar structures accompanied by structural changes of the repair condensates from fuzzy to round (Nagamalleswari et al, co-submitted). We now sought to compare ^T^E, ^T^S1E and ^T^4S2E expressing cells after different treatments (mock, ML792, CPT and ML792 & CPT) with respect to their co-localization with the nucleolar protein Nucleophosmin (NPM1). As shown in Fig. 5c and Extended Data Fig. 5a, we found all EME1-variants co-localizing with NPM1 in the same repair condensates under all conditions tested. We observed differences in the nucleoli number, shape and EME1 intensity as a function of treatment and EME1 variant. This is an interesting observation as the number of nucleoli is variable and an increase in numbers and size are common features of aggressive cancers. However, more extensive work is needed to understand how sumoylation and EME1 variants regulate the nature of nucleolar repair condensates.

For our study, we conclude that EME1 and both SUMO-mimetic variants co-localize with NPM1 in nucleolar repair condensates. More broadly, these findings support the idea that EME1 is sumoylated on DNA in the presence of specific DNA structures, leading to retention at DNA lesions in the nucleolar repair condensates.

### Removal of EME1 from DNA lesion via degradation and nuclear export

Our previous analysis showed that mono-sumoylation and poly-sumoylation have opposing functions for the stability and intracellular localization of EME1. While mono-sumoylation causes stabilization of EME1 and nuclear import/retention, poly-sumoylation appears to signal for degradation and export, as depicted in Fig. 6a.

**Fig. 6.**
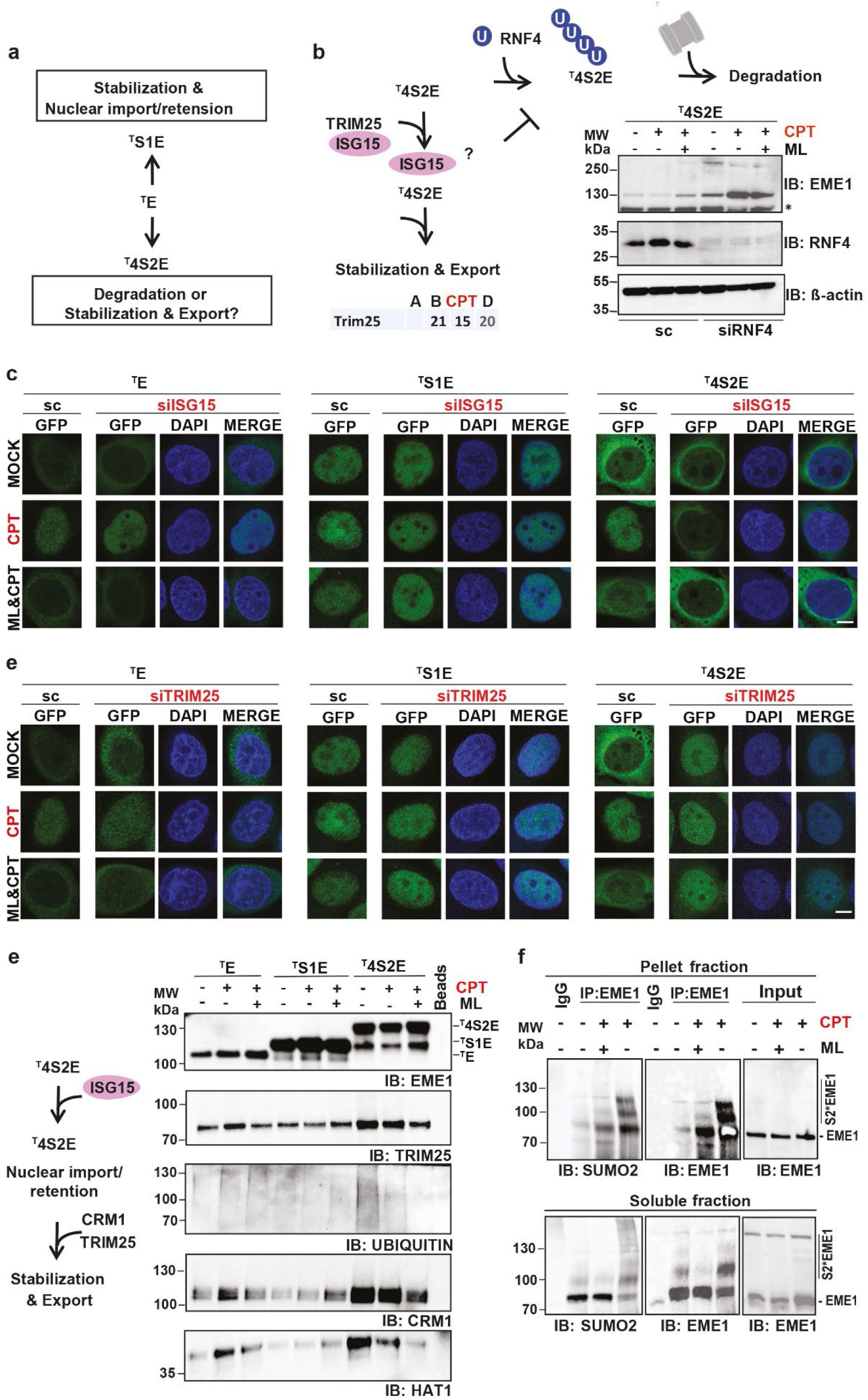
Removal of EME1 from DNA lesion via degradation and nuclear export & Endogenous EME1 is di- and poly-sumoylated upon CPT exposure. **(a)** Cartoon summary: EME1 mono-sumoylation (^T^S1E) causes stabilization and nuclear import or retention, while EME1 poly-sumoylation (^T^4S2E) might cause degradation or stabilization and nuclear export. **(b)** Cartoon working model: RNF4 dependent ^T^4S2E ubiquitination and degradation versus its ISG15 and TRIM25 dependent stabilization and nuclear export. Peptide numbers obtained from EME1 IP-MS analysis (Fig. 3a) for TRIM25. IgG (A) and EME1 immunoprecipitations (IP) upon mock (B), CPT (C) and CPT & ML792 (D) treatments. Immunoblot analysis of RNF4 depleted ^T^4S2E expressing cells. siRNA mediated knock down with scrambled (sc) or RNF4 (siRNF4) specific siRNAs for 48 hours before treatments DMSO (MOCK), CPT (10 µM for 1 hour) or ML & CPT (ML792, 1 µM for 2 hours). Cell lysates were resolved on SDS-PAGE and detected with anti-EME1 (^T^4S2E), anti-RNF4 and anti-β-actin antibodies. * indicates unspecific band. **(c)** Immunofluorescence analysis of ^T^E-variant (^T^E, ^T^S1E, and ^T^4S2E) expressing cell lines upon ISG15 depletion. siRNA mediated knock down with scrambled (sc) or ISG15 (siISG15) specific siRNAs for 48 hours before treatments with DMSO (MOCK), CPT (10 µM for 1 hour) or ML & CPT (ML792, 1 µM for 2 hours). Co-staining was with anti-GFP antibody (labelled with Alexa 488, green) for detecting ^T^E-variants and DAPI (blue) for visualization of nuclei. Shown are individual stains and merge of representative single cells. Scale bar represents 10 μm. Related multi cell images are shown in Extended Fig.6a and immunoblots confirming ISG15 knock down in Extended Data Fig. 6c. **(d)** Immunofluorescence analysis as **(c)** but upon TRIM25 depletion with TRIM25 specific siRNA (siTRIM25). Scale bar represents 10 μm. Related multi cell images are shown in Extended Data Fig. 6b and immunoblots confirming TRIM25 knock down in Extended Data Fig. 6d. **(e)** Carton working model: ISG15 regulates ^T^4S2E nuclear import/retention, while TRIM25 might result in stabilization and nuclear export. Immunoblots of GFP-Trap enriched ^T^E-variants and their interactions with indicated binding partners. ^T^E variant expression before treatments with DMSO, CPT or ML & CPT as in **(C)**. GFP-Trap-enriched samples were separated on SDS-PAGE and detected with anti-EME1, anti TRIM25, anti-Ubiquitin, anti-CRM1 and anti-HAT1 antibodies. Input control is shown in Extended Data Fig. 6e. **(f)** Immunoblots of EME1 immunoprecipitation (IP) upon DMSO, CPT or ML & CPT treatments as in **(c)** to detect endogenous EME1 sumoylation. Cells were lysed in RIPA buffer, sonicated and separated into the supernatant (soluble fraction) and pellet fractions. Pellets were resolved in RIPA buffer supplemented with 0.5 % SDS and after lysis diluted to 0.1 % SDS. Endogenous EME1 IPs were performed with anti-EME1 serum B antibodies. Samples were resolved on SDS-PAGE for supernatant and pellet fractions on different gels. Shown are immunoblots after detection with anti-EME1 serum A and anti-SUMO2 antibodies. Of note, EME1 migrates aberrant after the harsh treatment to resolve the pellet.

The SUMO-targeted ubiquitin E3 ligase RNF4 is best known for its function in ubiquitin dependent proteasomal degradation of poly-sumoylated substrates^41–43^. Hence, we tested its involvement in the degradation of the poly-sumoylated EME1 as outlined in Fig. 6b. RNAi-mediated downregulation of RNF4 in ^T^4S2E expressing cells after different treatments (mock, CPT and ML792 & CPT) showed that RNF4 is responsible for the degradation of ^T^4S2E in mock but to a greater extent in CPT treated cells (Fig. 6b). However, another mechanism must exist that protects a fraction of ^T^4S2E to allow nuclear export. The Chelbi-Alix lab recently proposed that TRIM25 dependent isgylation might protect poly-sumoylated substrates upon interferon response^44, 45^. Since exposure to CPT induces ISG15 expression and conjugation^46^, we revisited the EME1 interactome for putative candidates involved in the isgylation pathway. Indeed, we found TRIM25 in our screen (outlined in Fig. 6b). TRIM25 has been proposed as E3 ligase for ubiquitination and isgylation^47^. Hence, we studied whether RNAi mediated downregulation of ISG15 and TRIM25 had effects on the intracellular localization of ^T^E, ^T^S1E or ^T^4S2E after mock, CPT and ML792 & CPT treatments. Neither depletion of ISG15 (Fig. 6c and Extended Data Fig. 6a, c) nor TRIM25 (Fig. 6d, Extended Data Fig. 6b, d) had any effect on ^T^E and ^T^S1E, under all conditions tested. However, we observed opposite effects on ^T^4S2E: downregulation of ISG15 prevented nuclear import/retention of ^T^4S2E upon CPT exposure, whereas depletion of TRIM25 led to nuclear localization of ^T^4S2E, already in mock treated cells, and thus appeared to prevent its export. Interestingly, a recent study has shown that ISG15 is localized to replication forks and, when overexpressed, accelerates replication fork progression independent of conjugation, resulting in reduced CPT sensitivity^48^. Taken together, these findings support the idea that ISG15 may lock/retain ^T^4S2E at replication forks, although we cannot entirely exclude nuclear import functions. TRIM25 depletion (Fig. 6d) like CRM1 inhibition (Fig. 3c) block ^t^4S2E nuclear export, implying cooperative functions in the presence of SUMO chains, as depicted in Fig. 6e, left panel. To corroborate our model, we performed pull-downs to confirm interactions between transport receptor upon different treatments with the various EME1 variants (Fig. 6e and Extended Data 6e). Under the conditions used, we could not detect interactions with ISG15, IPO7, IPO9 and RNF4. It is likely that these interactions are transient or that the antibodies for detection are not sensitive enough. Though, in case of TRIM25, ubiquitin, CRM1 and HAT1, we observed by far the strongest interaction with ^T^4S2E in mock-treated cells, suggesting either efficient or very slow transport (Fig. 6e). The ^T^4S2E levels and interactions decreased upon CPT treatment, consistent with the RNF4 dependent degradation observed in Fig. 6b. ML792 & CPT stabilized ^T^4S2E but decreased all interactions, likely due to decreased nuclear import/retention and degradation that is dependent on additional sumoylation events. Intriguing, we detect highest ubiquitin levels, when RNF4 is least active, but these are concordant with TRIM25 levels, implying that TRIM25 might involve ubiquitination for its export functions. If this is true, TRIM25 would present a novel SUMO-targeted ubiquitin E3 ligase (STUbL), but this remains to be subjected to thorough biochemical analysis. Turning to ^T^E, we found that TRIM25, ubiquitin, CRM1 and HAT1 interact mainly upon CPT exposure in a ML792 reduced manner, suggesting that poly-sumoylation-dependent functions are enriched under this condition. ^T^S1E appears to be unregulated and shows little or no interaction despite its high abundance.

In summary, these data support that EME1 mono-sumoylation promotes nuclear import and retention, whereas poly-sumoylated might fosters nuclear import and export in addition to ubiquitin dependent degradation.

### Endogenous EME1 is di- and poly-sumoylated upon CPT exposure

Finally, we wanted to determine whether we could find evidence that endogenous EME1 is mono- and poly-sumoylated and performed large scale IPs from U2OS cell extracts that were mock, CPT or ML792 & CPT treated (Fig. 6f). Because EME1 variants were enriched in nucleolar repair condensates, we performed EME1 IPs from RIPA buffer soluble and the insoluble pellet fractions, latter were resolved at harsher conditions. Indeed, in the IP from the pellet fraction treated with CPT, we observed two slower migrating forms of EME1 that were reactive with SUMO2/3 and EME1 antibodies and were absent in extracts treated with ML792 & CPT, indicative for two mono-sumoylation or one di-sumoylation event (Fig. 6f). In the IP from the soluble RIPA buffer extracts, we detected a high molecular smear with SUMO2/3 antibodies, indicative for a SUMO chain (Fig. 6f). However, EME1 antibodies did not efficiently recognize this smear, probably because of limited sensitivity and epitope masking by the high modification levels.

In summary, these results demonstrate that endogenous EME1 becomes increasingly sumoylated upon CPT exposure, and enriches with two SUMO modifications in repair condensates and multiple-SUMO attachments in the soluble fraction. This is consistent with the notion that mono-sumoylation events are required for import and stabilization of EME1 at DNA lesions in nucleolar repair condensates and for subsequent poly-sumoylation dependent extraction from repair condensates for nuclear export or degradation.

## Discussion

The presented work sheds light on the multiple levels that control EME1, the regulatory subunit of the structure-specific endonuclease EME1-MUS81 complex. In our previous study, we observed that EME1 might be mono- and poly-sumoylated to promote CPT induced rDNA repair in the nucleolus (Nagamalleswari et al, co-submitted). Hence, we set out to understand the molecular consequences of these related but different modifications that appear to control EME1-MUS81 functions. We generated inducible cell lines expressing multiple EME1 variants: EME1 wild-type (^T^E), mono-sumo(1)ylation (^T^S1E) and poly-sumoylation (^T^4S2E) mimetic fusions, a mutant preventing sumoylation in close distance of EME1 (^T^UDE) and one that was mutated in its first out of two SIMs (^T^EΔSIM1). We aligned the main results with the endogenous protein and examined different regulators revealed by the endogenous EME1 interactome. Together, our data allow us to draw a first model how EME1 function is regulated upon replication stress as depict Extended Data Fig. 7.

Our analysis uncovered that EME1 is a shuttling protein that requires mono-sumoylation for its nuclear import and a functional SIM and poly-sumoylation for its nuclear export. Only marginal amounts of endogenous EME1 sumoylation are detectable at steady state, indicating that all sumoylation events are largely transient. However, large scale experiments (five 15 cm dishes/ lane demonstrated in Fig. 6f), suggested that sumoylation of endogenous EME1 is indeed enhanced upon CPT exposure, presumable with mono- and poly-sumoylation events. In unstressed cells, CRM1 and SIM dependent nuclear export predominates sumoylation dependent KPNB1 and IPO7 mediated import, placing EME1 mainly in the cytoplasm. Upon CPT exposure, the ratio turns in favor of its nuclear import, reinforced by HAT1 and acetylated H4 Histones that likely escort the EME1-MUS81 complex to sites of DNA replication. In the presence of stalled replication forks or nicked Holliday junctions resembling repair structures, EME1 is stabilized in repair condensates by a second mono-sumoylation event. Stabilization at these DNA structures promotes the induction of resolving “repair” DSBs, which act as prerequisite for replication. Subsequently, EME1 is poly-sumoylated, which facilitates DNA replication favored by ISG15 that might retain poly-sumoylated EME1 at DNA. Consistent with this, ISG15 is upregulated upon CPT exposure^46^ and has recently been shown to support DNA replication in a non-conjugated manner^48^. Then, poly-sumoylated EME1 is extracted by the AAA ATPase VCP/p97-NPL4-UFD1 complex, which we observed in our EME1-IP (Extended Data Fig. 7) and which is established to extract poly-sumoylated and/or ubiquitinated proteins form replication forks and nucleolar repair condensates^49–52^. This step likely requires poly-sumoylation-dependent ubiquitination, as it best understood for the STUbL RNF4 but may also apply to TRIM25. Depletion of RNF4 strongly stabilized ^T^4S2E in CPT-treated cells (Fig. 6b). Because we could not detect ^T^4S2E-RNF4 interaction and ubiquitination under this condition, we suspect transient interactions and efficient degradation. Though we detected ubiquitination concordant with ^T^4S2E-TRIM25 interaction (Fig. 6e) supporting the involvement of signaling ubiquitin functions. TRIM25 is indeed capable of assembling K63-linked ubiquitin chains like it best studied for RIG-I in context of activating the innate immune response^53^. It will be exciting to learn whether TRIM25 is a novel STUbL and whether DNA damage can directly trigger an immune response via this pathway from the nucleus to the cytoplasm.

An emerging question is how cells would decide between RNF4 mediated degradation and TRIM25 signaling pathways. One exciting candidate could be USP7, a SUMO-targeted ubiquitin protease that cooperates with VCP to define SUMO/ubiquitin levels at replication forks^50, 54^. In support, USP7 has been found to interact with TRIM25^55^, and has a clear preference for cleavage of K48- over K63-linked chains^56^. If co-recruited with TRIM25 it could prevent degradation in favor of signaling nuclear export.

Another question arising is which SUMO enzymes regulate the different EME1 sumoylation events. Our favored candidate for nuclear import dependent function is the nuclear pore associated RanBP2 as it binds to KPNB1^57^ and could modify EME1 on its transit to the nucleus^38^. Concordant, RanBP2 enhances EME1 sumoylation in *in vitro* sumoylation assays (Nagamalleswari et al, co-submitted) independent of DNA (Fig. 5a). Based on the co-localization with EME1 in nucleolar repair condensates, its preference for substrate mono-sumoylation and the enhanced activity in the presence of specific DNA structures, we propose the sumoylated UBC9 as the enzymes that executes the second EME1 mono-sumoylation event (Nagamalleswari et al, co-submitted). Our preferred candidates for EME1 poly-sumoylation are PIAS1 and PIAS4 as they assemble long SUMO chains on EME1 *in vitro* (Nagamalleswari et al, co-submitted), show enhanced modification in the presence of DNA (Fig. 5a) and are known to be recruited to CPT induced DNA lesions^58, 59^.

Next to the tight control of EME1, our data reinforce the important regulatory function of sumoylation in nucleo-cytoplasmic transport as it is discussed since its identification ^60–62^. We could not identify an export or import sequence in EME1, supporting piggy-backing or alternative mechanisms. When asked whether mono- and poly-sumoylation could act as import or export signals, we found in agreement that EME1, KPNB1, CRM1, TRIM25 and various nucleoporins can bind to SUMO^57, 63^. Since SUMO-tags improve protein solubility^64^, sumoylation could also be of great benefit for transport through the nuclear pore.

Export of EME1 depends on a functional SIM and CRM1, whereas its poly-sumoylated version additionally requires TRIM25. Both TRIM25 and CRM1 have been identified as low affinity interactors for di-SUMO^65^ and could therefore join forces in binding to poly-sumoylated EME1 for nuclear export. The fact that the SIM but not TRIM25 is required for export under mock conditions suggests involvement of intrinsic interactions with the attached SUMO chain, although we cannot exclude the involvement of other sumoylated proteins. However, we favor the idea that intrinsic SUMO chain-SIM interactions result in a closed conformation that is highly beneficial for transport through the nuclear pore. Since mono-sumoylated EME1 and the EME1 SIM1 mutant are trapped in the nucleus, we propose that di- or tri-sumoylated EME1 are required for intrinsic SIM binding to enable CRM1 interaction and export, while longer chains in addition depend on TRIM25. SUMO chain trimming might be performed by the SUMO chain editing protease SENP6. SENP6 is involved in degradation-independent functions, and EME1 was discovered as one of its substrates^66^. However, further functional and structural analyses are required to understand the underlying molecular mechanisms.

Regulatory subunits of MUS81 exist in two flavors, EME1 and the related, vertebrate-specific EME2, which shares 40% sequence identity to the C-terminus of EME1^67^. Although the S-phase specific EME2-MUS81 complexes account only about 20% of all MUS81 complexes, this complex appears to be the predominant complex responsible for DSB formation in S-phase, whereas EME1-MUS81 functions mainly in M-phase^2, 3, 68^. Our data suggest that EME1-MUS81 have specific functions in S-phase upon replication stress in repetitive DNA. The highly repetitive rDNA that is enriched in the nucleolus, the cellular compartment particularly prone to CPT induced DNA damage^69, 70^. Repetitive DNA is unique in its error-prone homologous recombination property and thus likely requires a specific repair mechanism to maintain genome stability (Nagamalleswari et al, co-submitted). Due to its repetitive nature and efficient transcription the nucleolus is subjected to constitutive replication stress, which explains why we observe survival, DSBs and replication are already increased in mock-treated cells. The regulatory mechanism we describe here is specific to EME1 as EME2 lacks the N-terminus that carries the major sumoylation site and the two SIMs (Nagamalleswari et al, co-submitted).

Overall, our findings on EME1-MUS81 regulation provide a key example of the multi-level monitoring of enzyme function to maintain genome stability in the stress-sensitive nucleolus. Since EME1 and MUS81 are among the top hits counteracting CPT-induced cell damage^12^, our results open up new approaches to interfere with EME1-MUS81 functions to reduce cancer drug resistance.

## Methods

### Antibodies

Rabbit polyclonal-antibodies were raised by Coring System Diagnostix GmbH against our purified recombinant human GST- and untagged EME1 full-length protein, validated (Nagamalleswari et al, co-submitted) and used as serum (serum A and B are from two rabbits). All other primary antibodies were purchased and validated by the manufacturers (data available on manufacturers’ websites).

#### Antibodies used in Immunoblot analysis (IB)

Primary antibodies used: rabbit anti-EME1 serum A & B (Coring System Diagnostix GmbH), mouse anti-RNF4 (MA527423, Life technologies), mouse anti-GFP (1814460, Roche), goat anti-IPO7 (PA518078, Invitrogen), rabbit anti-Histone H4 (acetyl K5 + K8 + K12 +K16 - ab177790, Abcam), rabbit anti-HAT1 (PA529174, Thermo fischer), mouse anti-ISG15 (PA579522, Invitrogen), mouse anti-TRIM25 (MA531936, Life technologies), mouse anti-Ubiquitin (sc-8017, Santa Cruz), rabbit anti- CRM1 (PA141634, Thermo Fisher) and mouse anti-β-actin antibody (A2228, Sigma-Aldrich). Mouse anti-SUMO2/3 (8A2) from DSHB deposited to the by Matunis. Secondary-antibodies used: donkey anti-rabbit (711-035-152, Jackson IR Ltd.), donkey anti-mouse (715-035-150, Jackson IR Ltd.), and donkey anti-goat antibody (705-035-003, Jackson IR Ltd.) horseradish-peroxidase conjugated.

#### Antibodies used in IF

Primary antibodies used: goat anti-GFP FITC (NB100-1771, Novus Biologicals), rabbit anti-EME1 serum A (Coring System Diagnostix GmbH), and mouse anti-NPM1 (Nucleophosmin 1) (B23, B0556, Sigma-Aldrich). Secondary antibodies used: anti-rabbit coupled to Alexa 594 dye (A32740, Invitrogen), anti-mouse coupled to Alexa 594 dye (715-585-150, Jackson IR Ltd.).

#### Antibodies for FACS analysis

anti-γΗ2ΑΧ coupled Alexa 647 (9720, Cell signaling), anti-GFP antibody FITC (NB100-1771, Novus Biologicals).

### Plasmids

For the inducible cell lines HA-Strep-GFP tagged full-length EME1-variants were cloned into pcDNA5: ^T^E, wild-type; ^T^S1E, SUMO1 fused N-terminally to EME1 representing mono-sumoylation; ^T^4S2E, linear fusion of four SUMO2 units (Eisenhardt et al, 2015) N-terminally to EME1 representing poly-sumoylation; ^T^UDE, catalytic domain of the yeast SUMO isopetidase Ulp1 ^1^ was fused N-terminally to EME1 to prevent sumoylation in close distance; ^T^EΔSIM, SIM1 amino acids 54 to 57 were mutated to alanins to prevent SUMO binding. MUS81 and EME1 were generated by PCR and cloned into pET23a+ and pET28a+, respectively. MUS81 and EME1 amplification from human cDNA. For antibody raising, EME1 was cloned into pGEX6P1. All constructs generated were verified by sequencing at the MPI-IE DNA Sequencing Core Facility.

### siRNA

Human siRNAs for the knock down assays were ordered from OriGene (IPO7: SR307155, HAT1: SR305590, ISG15: SR322849, TRIM25: SR305220, RNF4: SR304084, scrambled siRNA: SR30004).

### Cell lines & culture

U2OS (Human bone osteosarcoma epithelial cells) were grown in McCoy’s 5A (Modified) medium (16600082, Thermo Fischer) supplemented with 10% fetal bovine serum (FBS) (F0804, Sigma-Aldrich) and penicillin-streptomycin (P4458, Sigma-Aldrich). Hygromycin (CP12.2, Carl Roth), blasticidin (15205, Sigma-Aldrich) and Tet System approved FBS (631106, Takara) containing medium was used to culture doxycycline (Dox) (A2951, PanReac Applichem) induced stable cell lines. Subconfluent cultures were maintained at 37 °C in a humidified incubator with 5% CO_2_ and 95% air atmosphere.

U2OS cells modified with the Flp-In TRex system were a kind gift from Dr Jakob Nilsson. ^T^E, ^T^S1E, ^T^4S2E, ^T^EΔSIM, and ^T^UDE stable cell lines were generated in U2OS cells according to manufacturer’s instructions (Flp-In TRex manual, Invitrogen). Single clones were verified for dox inducible expression of the appropriate ^T^E, ^T^S1E, ^T^4S2E, ^T^EΔSIM, and ^T^UDE variants, via PCR, western blotting, and microscopy.

### Immunofluorescence assay (IFA)

^T^E, ^T^S1E, ^T^4S2E, ^T^EΔSIM, and ^T^UDE cell lines were seeded in Nunc Lab-Tek chambered slides and cells were either siRNA transfected (for 32 hours) and/or induced for 16 hours with Dox: ^T^E, ^T^S1E, ^T^ΔSIM - 4 ng/ml, ^T^4S2E - 10 ng/ml and ^T^UDE -1 µg/ml, respectively in Tet FBS medium. Subsequent cells were treated by DMSO (vehicle control), Camptothecin (CPT, C9911, Sigma-Aldrich) 10 µM for 1 hour, ML (ML792, gift from UbiQ) 1 µM for 2 hours) and ML & CPT (ML as added an hour before addition of CPT). For localization of endogenous EME1, U2OS cells were seeded and treated in the same manner as the other cell lines. After treatments, cells were washed with 1X PBS (14190250, Thermo Fischer), followed by fixation with 4% formaldehyde (F8775, Sigma-Aldrich) for 15 minutes and permeabilization with 0.5% triton X 100 (3051, Carl Roth) for 10 minutes. Coverslips were then blocked with Sea-block buffer (37527, Thermo Fischer) for 1 hour at room temperature, washed with 0.2% triton X 100 and incubated with primary antibodies (1:300 dilution in Sea-block buffer) overnight at 4°C. Further washing with 0.2% triton X 100, followed by a one-hour incubation with appropriate secondary antibodies at room temperature (1:1000 dilution in Sea-block buffer). The slides were washed again before being fixed with anti-fade DAPI (S36939, Thermo Fischer). Primary antibodies used were anti-GFP FITC (NB100-1771, Novus Biologicals), or anti-EME1 serum A (Coring System Diagnostix GmbH) for detecting endogenous EME1. Secondary antibody for detecting EME1 was anti-rabbit coupled to Alexa 594 dye (A32740, Invitrogen). Images were acquired with a Zeiss LSM780 confocal microscope using 63x oil immersion objective at a zoom range 0.6-2x. Images were processed using Zen black & blue or Imaris software.

For Leptomycin B (LMB) (9676S, Cell signaling) treatment, induced cells were treated with ethanol (MOCK), and 80 nM LMB for 3 hours, followed by Immunofluorescence assay.

For Importazole (IPZ) (SML0340, Sigma-Aldrich) treatment, induced cells were treated with DMSO (MOCK), and 75 µM IPZ for 2 hours, followed by 1 hour treatment with CPT. Above mentioned Immunofluorescence assay was carried out.

For Trichostatin A (TSA) (T8552, Sigma-Aldrich) treatment, induced cells were treated with DMSO (MOCK), and 600 nM TSA for 16 hours, followed by treatment with DMSO (MOCK), ML, CPT and ML & CPT. Above mentioned Immunofluorescence assay was carried out.

### Analysis of nuclear/nucleolar condensates by pre-extraction with CSK-buffer

Pre-extraction to release soluble proteins and enrich in nuclear condensates was performed by pre-incubation with CSK buffer as described in^2, 3^. In brief, after seeding and treatments of cells as described above, coverslips were washed and incubated with CSK buffer (10 mM PIPES with pH 7.0, 100 mM NaCl, 300 mM sucrose, 3 mM MgCl2 and 0.7% Triton X-100) for 3 minutes at room temperature. Subsequently, cells were washed, fixed with 4% paraformaldehyde (PFA) (SC281692, Santa Cruz) in PBS for 25 minutes at room temperature before staining the cells with the respective antibodies as described above. For nucleolar staining, cells were incubated with anti-NPM1 (Nucleophosmin 1) (B23, B0556, Sigma-Aldrich). Secondary antibody for detecting NPM1 was anti-mouse coupled to Alexa 594 dye (715-585-150, Jackson IR Ltd.). Slides were fixed using anti-fade DAPI.

### RNAi transfection

^T^E, ^T^S1E, and ^T^4S2E cells were seeded and IPO7, HAT1, ISG15, TRIM25, and RNF4 knockdown was performed using their respective siRNA with Lipofectamine RNAiMAX transfection reagent (13778-075, Invitrogen) as per manufacturer’s instructions for 48 hours. 32 hours post RNAi transfection, ^T^E, ^T^S1E, and ^T^4S2E cells were induced by Dox (4, 4, 10 ng/ml respectively) followed by treatments. Further performed Immunofluorescence assay, and Immunoblotting.

### Flow cytometry

^T^E, ^T^S1E, ^T^4S2E, ^T^EΔSIM and ^T^UDE expressing tagged cell lines were seeded in 10 cm dishes and induced for 16 hours with Dox: ^T^E, ^T^S1E, ^T^EΔSIM - 4 ng/ml, ^T^4S2E - 10 ng/ml and ^T^UDE -1 µg/ml respectively. Cells were treated with 1 µM ML792 for 2 hours and 10 µM CPT for 1 hour, followed by EdU pulse (10 µM) for 30 minutes. Trypsinized cells were washed with 1% BSA in PBS, and fixed with 4% paraformaldehyde (PFA) for 15 minutes at room temperature. After additional washing, cells were permeabilized with the 1X Saponin based solution provided in Click iT Kit (C10646, Thermo Fischer) for 15 minutes and treated with Click-iT Plus EdU reaction mix as per manufacturer’s instructions (Click-iT Plus EdU Alexa Fluor 594 Flow Cytometry, C10646, Thermo Fischer). Samples were incubated at room temperature for 30 minutes, washed with 1X Saponin based wash buffer, blocked with Sea-block buffer (37527- Thermo Fischer) for one hour at room temperature and co-stained with Alexa 647-coupled mouse anti-γH2AX (560447, BD Biosciences) and FITC coupled goat anti-GFP (NB100-1771, Novus Biologicals) antibodies. DAPI (D1306, Thermo Fischer) was added to a final concentration of 1 mg/ml to facilitate exclusion of dead cells from the analysis and cells were resuspended in 500 µl of cold PBS. Samples were analyzed with a Fortessa III flow cytometer (BD Biosciences) and FlowJo software. Graphs were plotted using Graphpad Prism 8.0.

### Laser microirradiation

Laser track experiments were performed using a Leica SP5 confocal microscope with an environmental chamber set to 37 °C. Briefly, ^T^E, ^T^S1E, and ^T^4S2E cell lines were grown on 18 mm coverslips and media was replaced by fresh medium with 20 µg/mL Dox. Prior to micro irradiation, medium was replaced by CO2-independent Leibovitz’s L15 medium supplemented with 10% FCS, Penicillin/Streptomycin and 20 µg/mL Dox. Laser micro-irradiation was carried out on a Leica SP5 confocal microscope equipped with an environmental chamber set to 37°C. DNA damage tracks (1 µm width) were generated with a Mira modelocked titanium-sapphire (Ti:Sapphire) laser (l = 800 nm, pulse length = 200 fs, repetition rate = 76 MHz, output power = 80 mW) using a UV-transmitting 63 × 1.4 NA oil immersion objective (HCX PL APO; Leica). Gain was set at 30 % and offset at 50 %, output was 1.5 ± 0.5 W. Relative recruitment values were analyzed using Leica LAS X Software.

### Clonogenic survival assays

^T^E, ^T^S1E, and ^T^4S2E cell lines were seeded in 6 well plate and induced with Dox (4, 4, and 10 ng/ml respectively) for 16 hours. Cells were treated with CPT (0, 10, 100 and 500 nM) for 1 hour, DMSO was used as a control. Subsequently, cells were washed with 1X PBS, trypsinized and seeded in triplicates into 6-well plates at a density of 500 cells per well. After 5-7 days post CPT treatment, the cells were washed, fixed with PFA and stained with 0.5% Crystal Violet solution (in 20% methanol) and photographed on a light table. The images were analyzed with ImageJ and measure absorbance at 595 nm. Plots and statistics were performed using Graphpad Prism 8.0.

### Cell extracts for Immunoblotting (IB)

Dox induced cells were harvested and lysed in lysis buffer to proceed for Western blotting (50 mM Tris-Cl pH-6.8, 0.1% bromophenol blue, 10% glycerol, 100 mM DTT, 5 mM EDTA, 2% SDS). Lysed protein samples were sonicated and resolved by SDS-PAGE. Proteins were transferred onto the nitrocellulose membrane using semi-dry transfer apparatus. Blots were blocked using PBST (1X PBS + 0.1% Triton X 100) with 5% skimmed milk. Eventually, the blots were probed with indicated primary antibodies: anti-EME1 serum A (homemade), anti-RNF4 (MA527423, Life technologies), anti-GFP (1814460, Roche), anti-IPO7 (PA518078, Invitrogen), anti-HAT1 (PA529174, Thermo fischer), anti-ISG15 (PA579522, Invitrogen), anti-TRIM25 (MA531936, Life technologies) and anti-β-actin antibody (A2228, Sigma-Aldrich), incubated overnight at 4°C at 1:2000 dilution. Further incubated with secondary antibodies: anti-mouse (715-035-150, Jackson IR Ltd.), anti-rabbit (711-035-152, Jackson IR Ltd.) and anti-goat antibody (705-035-003, Jackson IR Ltd.) HRP at 1:5000 dilution for 1 hour at room temperature. Detection was with ECL reagent (1705061, Biorad).

### Immunoprecipitation (IP)

U2OS cells were seeded in 4 x 15 cm dishes and cells were treated with 1 µM ML792 for 2 hours, 10 µM CPT for 1 hour and PR619 for 30 minutes (PR619 blocks ubiquitin and SUMO proteases-16276, Cayman Chemical). Cells were washed, scraped and lysed in a RIPA buffer (20 mM Tris pH 7.5, 150 mM NaCl, 5 mM EDTA, 1% NP40, 5% glycerol, 0.5% Na-deoxycholate, 0.1% SDS) supplemented with protease inhibitors [1 mM Leupeptin (A2183 , Merck ), Pepstatin A (A2205, Merck), 1 mM Aprotinin (A1623, Roth), 100 mM Pefabloc (11873601001, Merck), 400 mM Iodoacetamide (I1149, Sigma-Aldrich), 400 mM N-Ethylmaleimide (E1271, Sigma-Aldrich) and 200 mM Orthovanadate (P07581, New England Biolabs)]. Lysed cells were sonicated, and then separated in supernatant (soluble fraction) and pellet fractions. Pellets were dissolved in RIPA buffer containing 0.5 % SDS, then lysed, additional sonication before further dilution with dilution buffer (20 mM Tris pH 7.5 and 150 mM NaCl along with protease inhibitors as mentioned above) to reduce SDS concentration from 0.5 % to 0.1 % as it is required for IPs (pellet fraction).

Subsequent immunoprecipitations were performed from both fractions by incubating lysates with anti-EME1 serum B polyclonal antibodies (homemade) overnight at 4°C. Immunoprecipitates were collected on pre-equilibrated (RIPA buffer) protein A Sepharose beads (53139, Thermo Scientific) by incubating the reaction for 2 hours at room temperature on a rotating wheel. After additional washing steps with RIPA buffer, beads were boiled in 2X Laemmli buffer for 5 minutes at 95°C. Samples were resolved by 7% SDS-PAGE and transferred to nitrocellulose membrane by semi-dry i-blot transfer system (IB1001, Invitrogen). Inputs and elution fractions were analyzed on separate gels. Blots were incubated with primary antibodies: EME1 serum A (homemade), anti-Histone H4 (acetyl K5 + K8 + K12 +K16 - ab177790, Abcam), anti-HAT1 (PA529174-Thermo Fisher Scientific) or anti-SUMO2 (DSHB) antibodies at 1:2000 dilutions, overnight at 4°C, followed by incubation with anti-rabbit secondary antibody (711-035-152, Jackson IR Ltd.) at 1:10.000 dilution for 1 hour at room temperature. Detection was with ECL reagent (1705061, Bio-Rad) on a Bio-Rad ChemiDoc imaging system (17001402, Bio-Rad).

### Mass Spectrometric analysis for endogenous EME1-IP

#### MS sample preparation

IP samples were eluted with 40 µl 1xLDS buffer adjusted to 10 mM DTT (5 min, 90°C) and separated on 4-12% Bis-Tris NuPAGE gels (Thermo Fisher) followed by colloidal coomassie staining (Instant Blue, Expedeon). Each lane was sliced into nine gel bands and the excised gel pieces were subjected to in-gel digestion using trypsin (Promega) followed by C18 STAGE tip clean-up (as described in ^4^).

#### NanoLC-MS analyses

Nano-scale LC-MS analysis was done on a Q Exactive mass spectrometer coupled to an EASY-nLC 1000 nUHPLC (both Thermo Fisher Scientific) essentially as described in ^5^ with modifications detailed below. Peptide samples (reconstituted in 10 µl at 0.15% trifluoroacetic acid, 0.09% formic acid, 0.75% acetonitrile) were injected (3 µl) twice (60 min nLC-MS method). The gradient was: 5 min: 10%, 40 min: 40%, 4 min: 80% (250 nl/min flow rate). This was followed by a “wash out step”: 5 min: 80%B buffer and a 5 min inverse gradient from 80% to 2% B buffer (flow rates 450 nl/min). All measurements were carried out in data-dependent mode employing the “sensitive method” ^5^.

#### Mass spectrometry data analysis

MaxQuant (v. 1.6.5.0; ^6^) employing standard parameters (except minimal peptide length was set to six and MS/MS tol. (FTMS) was set to 25 ppm) was utilized to identify peptides and pursue final protein identification role-up (both at 1%FDR). MS raw data were searched simultaneously with the target-decoy standard settings against the Uniprot Homo sapiens database (Uniprot_reviewed+Trembl including canonical isoforms; downloaded on January 10^th^, 2017) and an in-house curated FASTA file (containing commonly observed contaminant proteins). Trypsin/P was used as enzyme and up to 2 missed cleavages were allowed. Carbamidomethylation of cysteine was set as fixed modification. Variable modifications included oxidation (M), deamidation (N, Q) and acetylation of protein N-termini. The match-between-run (MBR) option was disabled and intra sample abundance was approximated by intensity-based absolute quantification (iBAQ). For each of the four conditions (IP IgG MOCK, IP EME1 MOCK, IP EME1 CPT and IP EME1 ML&CPT), the nine gel slices were labeled with a unique “fraction number” in MaxQuant, respectively. The latter enabled the inter-sample comparison of individual peptide counts for a given protein of interest on basis of corresponding gel slices (Fig. 3B).

### Mass spectrometry analysis for the His-SUMO pull downs upon TSA treatment

#### Sample preparation

His10-SUMO2 conjugates were enriched as described previously ^7^. Briefly, Hela parental and His10-SUMO2 expressing cells were treated with DMSO or histone deacetylase inhibitor (HDACi) trichostatin A (TSA) 600 nM for 18 hours. Post treatment the cells were washed thrice, scraped, and collected in ice-cold PBS. A small aliquot of cells was separately lysed in 2% SDS, 1% N-P40, 50mM TRIS pH 7.5 and 150mM NaCl for total lysates. 6M guanidine-HCl pH 8.0 (6 M guanidine-HCl, 0.1M Na2 HPO4 /NaH2 PO4, 10mM TRIS, pH 8.0) was used to lyse the remaining parts of the cell pellets. Lysates were sonication twice for 5 seconds, using a sonicator (Misonix Sonicator 3000, EW-04711–81) at 30W to homogenise lysates. Protein concentrations were determined using the bicinchoninic acid (BCA) protein assay reagent (Thermo Fischer) and lysates were equalised. 50mM final concentration of imidazole was added to each lysates. Further these lysates were incubated overnight at 4°C with prewashed Ni-NTA beads (Qiagen, 30210).Sequential washes with wash buffer 1-4; Wash buffer 1: 6M Guanidine-HCL, 100mM sodium phosphate, 10mM Tris, 10mM imidazole, 5mM β-mercaptoethanol, 0.2% Triton X-100. Wash buffer 2: 8M urea, 100mM sodium phosphate, 10mM Tris, 10mM imidazole, 5mM β-mercaptoethanol, 0.1% Triton X-100. Wash buffer 3: 8M urea, 100mM sodium phosphate, 10mM Tris, 10mM imidazole, 5mM β-mercaptoethanol. Wash buffer 4: 8M urea, 100mM sodium phosphate, 10mM Tris, 5mM β-mercaptoethanol, were carried out to reduce background binding. Enriched sumo conjugates were eluted twice in one bead volume of 7M urea, 100mM sodium phosphate, 10mM Tris and 500mM imidazole pH 7.0.

#### Proteomic sample preparation

Pre-washed 100 kDa cut off filters (Elution Buffer minus imidazole) were used to concentrate His10 SUMO2 enriched fraction. The concentrates were diluted with ammonium bicarbonate to an end concentration of 50mM. Reduction and alkylation was carried out by adding DTT to 5mM and chloroacetamide to 5mM subsequently. All the steps were carried out at room temperature for 30 minutes each. The samples were diluted by adding 3 volumes of 50mM ammonium bicarbonate and trypsin digestion was carried out at a 1:50 enzyme-to-protein ratio, overnight and in the dark at room temperature. The digested peptides were acidified with 2% TFA and desalted and concentrated on triple-disc C18 reversed phase StageTips. Peptides were eluted with acetonitrile (ACN), vacuum dried and dissolved in 0.1% formic acid (FA) prior to liquid chromatography-tandem mass spectrometry.

#### Mass spectrometry

Samples were run on an EASY-nLC 1000 system (Proxeon, Odense, Denmark) connected to a Q-Exactive Orbitrap (Thermo Fisher Scientific, Schwerte, Germany) through a nano-electrospray ion source. Peptides were separated in a 15 cm analytical column (MS Wil, Aarle-Rixtel, The Netherlands) with an inner-diameter of 75 μm, in-house packed with 1.8 μm C18 beads (Reprospher-DE, Pur, Dr. Manish, Ammerbuch-Entringen, Germany). The gradient was 120 min from 2% to 95% acetonitrile in 0.1% formic acid at a flow rate of 200 nL/minute. Full-scan MS spectra were acquired with an Automatic Gain Control (AGC) target of 3e6 and a resolution of 70,000, maximum injection time of 20ms and a scan range of 400 to 2000 m/z. The top10 most abundant ions were sent for MS2 at a resolution of 17,500, AGC target of 1e5, maximum injection time of 60ms, a loop count of 10, dynamic exclusion set to 60ms and a scan range of 400 to 2000 m/z. The higher-collisional dissociation (HCD) normalized collision energy (NCE) of 25%. Ions with charge 1, and greater than 6 were excluded from triggering MS2 events. Each sample was injected in two technical repeats.

#### MS raw data processing

MS data files were analysed using MaxQuant (version 1.6.5) matching to Uniprot Human Proteome (Reviewed human proteome downloaded from Uniprot on 4^th^ of October 2019) with default settings of FDR and andromeda score filtering, matching to decoy database and common contaminants. Digestion was set to allow four missed cleavages with Trypsin digestion. Normalization was done by LFQ (default settings) with matching between runs and matching unidentified features enabled. Cysteine carbamidomethylation was set as fixed modification and N-terminal acetylation, oxidation of methionine were included as variable modifications.

#### Bioinformatic data analysis of MS data

Analysis was performed in Perseus (version 1.6.7) on ProteinGroups.txt output file. Proteins only identified by site, hits to the decoy database and potential contaminants were filtered out. Values were log2 transformed and filtered to include at least three valid values in one treatment group, remaining missing values were imputed from the lower end of the normal distribution (width: 0.3, down shift: 1.8). Background binders were excluded by performing a right-sided student’s t-test (S0=0.1, FDR=0.05) between parental samples and His10-SUMO2 expressing cells, only proteins significantly enriched in His10-SUMO2 were kept. Next, a two-sided student’s t-test between His10-SUMO2 expressing cells treated with TSA or DMSO was performed and plotted as a volcano plot.

### Pull downs by GFP-Trap

HA-Strep-GFP tagged ^T^E, ^T^S1E and ^T^4S2E cell lines were grown in 1 X 15 cm dishes and induced with Dox for 16 hours: ^T^E, ^T^S1E - 200 ng/ml and ^T^4S2E - 1ug/ml respectively. Cells were treated with DMSO as control, 1 µM ML792 for 2 hours, 10 µM CPT for 1 hour and PR619 for 30 minutes (PR619 blocks ubiquitin - 16276, Cayman Chemical). Cells were lysed in lysis buffer (10 mM Tris pH 7.5, 150 mM NaCl, 0.5 mM EDTA, 1% Triton X 100, 5% glycerol, 1% Na-deoxycholate, 0.1% SDS and supplemented with protease inhibitors [1 mM Leupeptin (A2183, Merck), Pepstatin A (A2205, Merck), 1 mM Aprotinin (A1623, Roth), 100 mM Pefabloc (11873601001, Merck), 400 mM Iodoacetamide (I1149, Sigma-Aldrich), 400 mM N-Ethylmaleimide (E1271, Sigma-Aldrich) and 200 mM Orthovanadate (P07581, New England Biolabs)] and incubated at 4°C on a rotating wheel for 30 minutes. Samples were sonicated and centrifuged then diluted with dilution buffer (10 mM Tris PH 7.5, 150 mM NaCl, 0.5 mM EDTA). Lysates were pre equilibrated with dilution buffer and incubated with GFP Trap beads (gta-10, Chromotek) on a rotating wheel at 4°C overnight. Beads were washed with dilution buffer and samples were eluted with 2X Laemmli buffer for 5 minutes at 95°C. Samples were resolved on a 4-12 % gradient SDS-PAGE (NW04122, Thermo Fisher) and transferred to nitrocellulose membrane by semi-dry i-blot transfer system (IB1001, Invitrogen). Blots were incubated with primary antibodies: anti-EME1 rabbit serum A (Coring system Diagnostix GmbH), anti-MUS81 (M1445, Sigma-Aldrich), anti-TRIM25 (MA531936, Life Technologies), anti-Ubiquitin (sc-8017, Santa Cruz), anti-CRM1 (PA141634, Thermo Fisher) and anti-HAT1 antibody (PA529174, Thermo Fisher). Further incubated with anti-mouse (715-035-150, Jackson), anti-rabbit secondary antibody (711-035-152, Jackson IR Ltd) HRP at 1:10,000 dilution for 1 hour at room temperature. Detection was with ECL reagent (1705061, Biorad) on a Bio-Rad ChemiDoc imaging system (17001402, Bio-Rad).

### Protein purification

All bacterial expression constructs were transformed into chemically competent bacteria, cultured in lysogeny broth (LB) and selected with the appropriate antibiotics.

Purification of mammalian SUMO2, SUMO E1 (AOS1/UBA2), SUMO E2 (UBC= WT), GST-RanBP2ΔFG, 6xHis-PIAS1, 6xHis-MPB-ZNF451-N, 6xHis-MPB-ZNF451-3 and S2*Ubc9 has been described^8–11^. MUS81-6x-His-EME1 complex was purified as described (Nagamalleswari et al., co submitted).

### *In vitro* sumoylation assay

The assays were performed as described before^8–11^. Briefly, reactions were performed in 20 μl volumes containing 20 mM HEPES pH 7.3, 110 mM potassium acetate, 2 mM magnesium acetate, 0.05% (v/v) Tween20, 1 mM DTT, 0.2 mg/ml ovalbumin, and 5 mM adenosine triphosphate (ATP). The DNA substrates used for the assays were prepared as previously described ^12, 13^. Different concentrations of DNA ( 4, 20 and 100 nM) - replication fork (RF), Holliday junction (HJ) single stranded DNA, double stranded DNA, and plasmid were used in the assay. Samples were incubated at 30°C for 30 minutes. Reactions were terminated by boiling with 1X Laemmli buffer at 95°C for 5 minutes and samples were resolved by SDS-PAGE. Detection was done as indicated.

### Endonuclease assays

The DNA substrates used for the endonuclease assays were prepared as previously described ^12, 13^. Instead of a 5’-32P-end label, we used a 5’-DY782-end-label on one oligonucleotide, which can be monitored using a Odyssey CL imager (LI-COR). Labelled oligonucleotides were obtained from Eurofins. Reactions were performed as specified in the Fig. legends. In brief, reactions were incubated for 1 hour at 37 °C and then stopped by addition of SDS and proteinase K to a final concentration of 0.8 % and 2 mg/ml respectively.

Subsequently, the reactions were incubated for another 15 minutes at 37 °C. Samples were mixed with 6 x loading buffer (30 % glycerol, 0.25 % bromphenol blue, 0.25 % xylene cyanol FF) and loaded on a 8 % native PAGE. After electrophoresis, the gels were scanned with an Odyssey CL imager (LI-COR).

### Statistics

Graphpad Prism v8.0 (Graphpad Software) was used to plot and analyze data. All experiments with quantitative analysis included data from at least 3 independent experiments. Data were expressed as mean ± SEM, and statistical differences were calculated by two-way ANOVA using Tukey test for grouped samples.

## Reporting summary

Further information on research design is available in the Nature Research Reporting Summary.

## Data availability

No source data are provided with this manuscript.

## Acknowledgements

We would like to thank all former and current members of A.P.’s laboratory for discussions and sharing reagents, and the MPI-IE core facilities for Proteomics, Imaging, and Flow cytometry for their technical help. ML792 was a kind gift from Farid El Oualid and Alfred Nijkerk (UbiQ, Amsterdam). For sharing reagents, we kindly acknowledge Jakob Nilsson. The Pichler lab is funded by the Max Planck Society. A.P. and A.C.O.V received funding from the European Union’s Horizon 2020 research and innovation program under the Marie Sklodowska-Curie grant agreement 765445. Román González Prieto was supported by the Dutch Cancer Society, (KWF-YIG 11367 / 2017-2). R.T. is supported EU Marie-Curie Leading Fellowship (number 707404). This article is based on work from European Cooperation in Science and Technology (COST) Action (PROTEOSTASIS BM1307 and ProteoCure CA20113 to A.P. and A.C.O.V.) and, supported by COST.

## Author contribution

J.B. and A.P. initiated and A.P. has set the direction of this study. K.B, E.N., J.B., N.E., R.G.P and A.P. designed the majority of experiments. H.L., N.E., and C.F. generated the stable EME1 variant inducible cell lines. All IFAs and clonogenic survival assays were performed by K.B.; J.B. expressed recombinant EME1 and MUS81 and performed nuclease assays and generated EME1 antibodies ; FACS analysis was carried out by E.N.; endogenous EME1 IP was conducted by N.E and E.N; and MS analysis by G.M.; TSA treatment was performed by R.T. and subsequent MS by F.T; RNAi treatments and pull downs from stable cell lines were carried out by E.N and K.B.; V.K.C. purified multiple proteins for in vitro sumoylation assays; endogenous EME1 sumoylation by E.N.; DNA dependent in vitro sumoylation assays were performed by N.E.; recruitment assays by R.G.P; statistical analysis by E.N., K.B., R.G.P.; D.I. helped with different experiments. A.P. supervised. K.B., E.N., J.B., C.F., N.E., D.I., R.G.P and A.C.O.V supervised. R.T, F.T, and R.G.P.; K.B., E.N., J.B., C.F., N.E., G.M., R.G.P., and R.T. prepared figures, figure legends and wrote the methods part. A.P. wrote the manuscript with the input from all authors.

---

**Extended Data Fig. 1.**
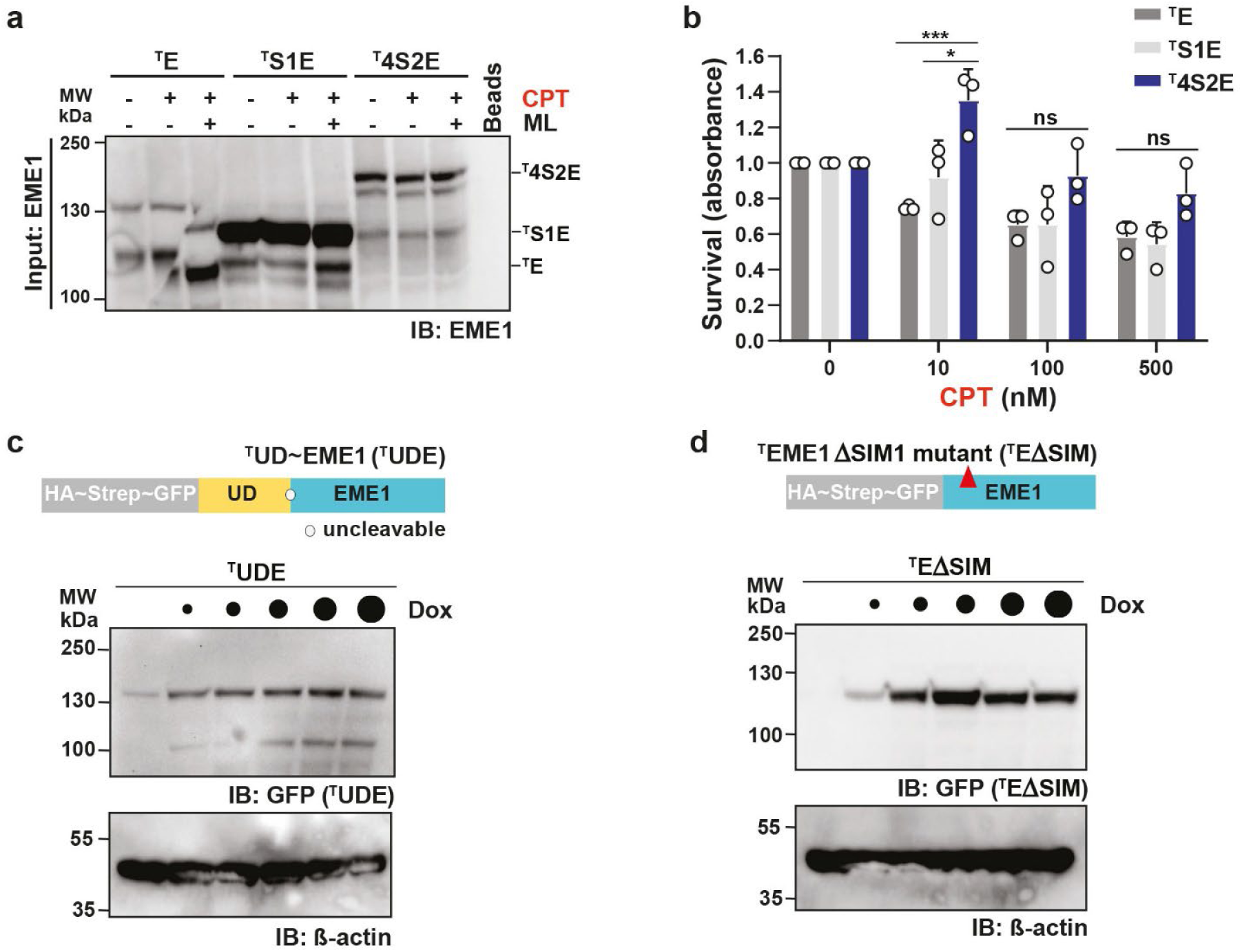
EME1 mono- and poly-sumoylation promote cell survival, DSBs & replication upon CPT exposure. Related to Fig. 1. **(a)** Input control to Fig. 1b. **(b)** As Fig. 1c but bar chart normalized to ^T^EME1 variant’s mock. **(c)** Schematic representation of the HA-Strep-GFP tagged EME1-variant ^T^UDE, in which the catalytic domain of the yeast SUMO isopetidase Ulp1 was N-terminally fused to EME1 to prevent sumoylation. Immunoblot shows expression levels of the stably integrated ^T^UDE induced by different Dox concentrations (0, 4, 20, 100, 1000 ng/ml) for 16 hours. Cell lysates were resolved on SDS-PAGE and ^T^UDE expression (anti-GFP) was monitored in comparison to anti-β-actin levels. **(d)** Schematic representation of the HA-Strep-GFP tagged EME1-variant ^T^EΔSIM, that is mutated in its first SUMO interaction motif (SIM). Immunoblot shows expression levels of the stably integrated ^T^EΔSIM induced by different Dox concentrations (0, 4, 20, 100, 1000 ng/ml) for 16 hours. Cell lysates were resolved on SDS-PAGE and ^T^UDE expression (anti-GFP) was monitored in comparison to anti-β-actin levels.

**Extended Data Fig. 2.**
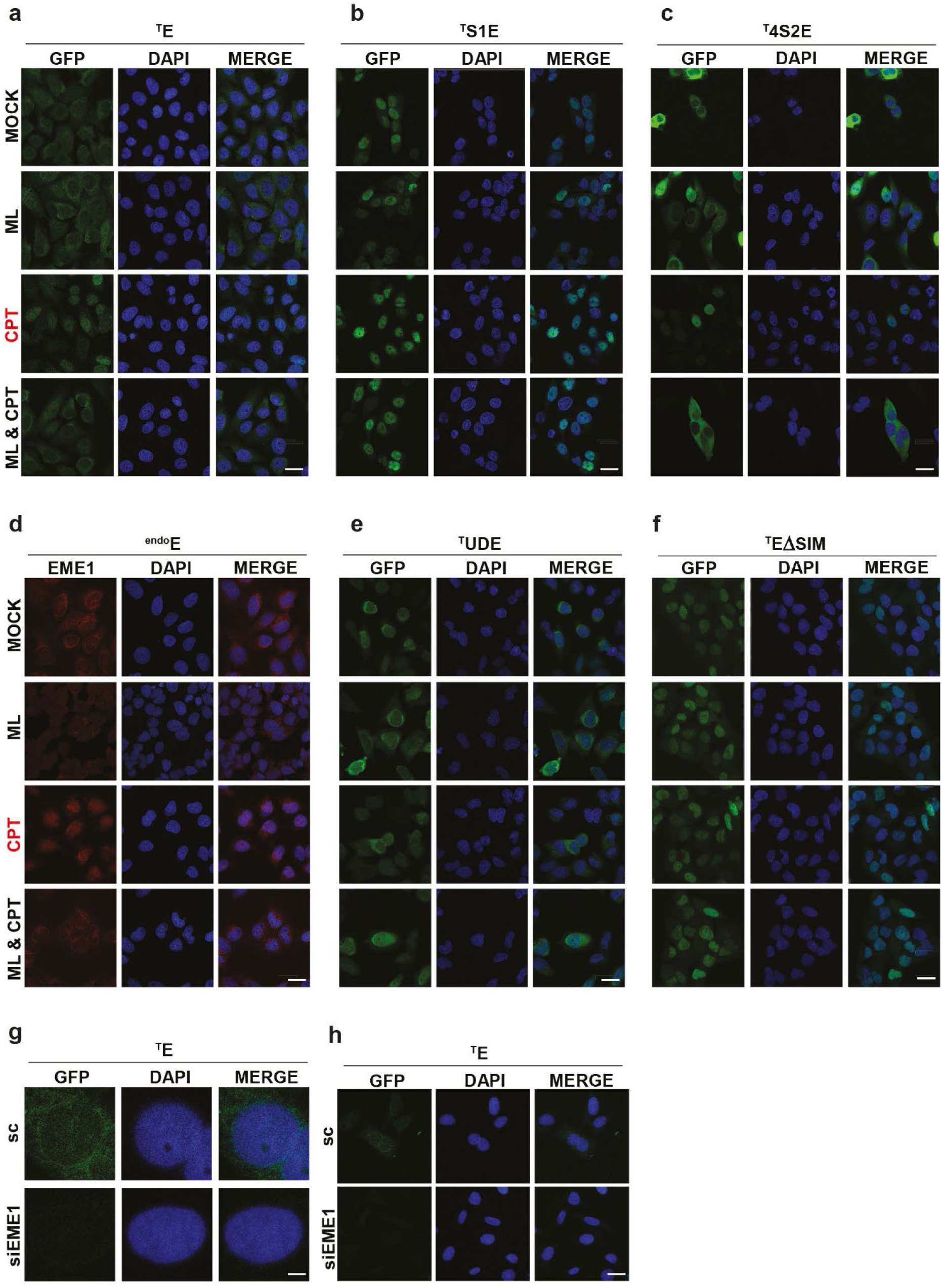
Sumoylation signature and SUMO/SIM interaction direct EME1’s intracellular placements. Related to Fig. 2. **(a)** Multi-cell images related to Fig. 2a. **(b)** Multi-cell images related to Fig. 2b. **(c)** Multi-cell images related to Fig. 2c. **(d)** Multi-cell images related to Fig. 2d. **(e)** Multi-cell images related to Fig. 2e. **(f)** Multi-cell images related to Fig. 2f. **(g)** Immunofluorescence analysis upon RNAi mediated knock down of EME1 for 48 hours in ^T^E expressing U2OS cells. 32 hours post siRNA transfection, cell line was induced by 4 ng/ml Dox concentration. Co-staining was with anti-GFP antibody (labelled with Alexa 488, green) for detecting ^T^E-variant and DAPI (blue) for visualization of nuclei. Shown are individual stains and merge of representative single cells. Scale bar represents 10 μm. **(h)** Multi-cell images related to (g).

**Extended Data Fig. 3.**
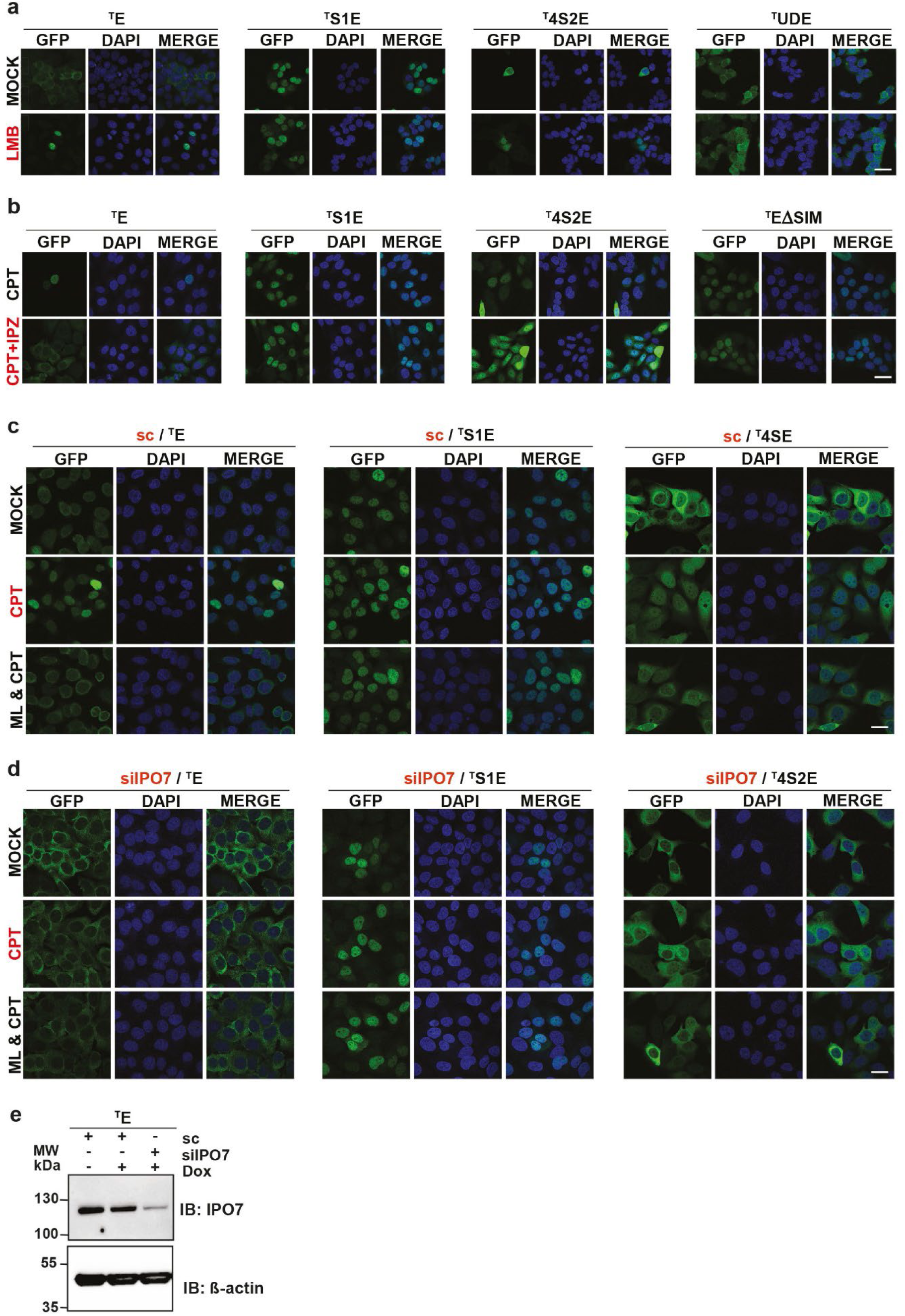
Import and export receptors involved in EME1’s intracellular localization. Multi-cell images and RNAi controls related to Fig. 3. **(a)** Multi-cell images related to Fig. 3c. **(b)** Multi-cell images related to Fig. 3d. **(c)** Full multi-cell image panels of ^T^E-variants upon control knockdown using scrambled siRNA (sc), related to Fig. 3e. **(d)** Full multi-cell image panels of ^T^E-variants upon IPO7 knockdown using siRNA, related to Fig. 3e. **(e)** Immunoblot verifying RNAi mediated knock down of IPO7 for 48 hours in ^T^E expressing U2OS cells. Related to Fig. 3e. 32 hours post siRNA transfection, cell line was induced by 4 ng/ml Dox concentration. Cell lysates were resolved on SDS-PAGE and IPO7 expression (anti-IPO7) was monitored in comparison to anti-β-actin levels.

**Extended Data Fig. 4.**
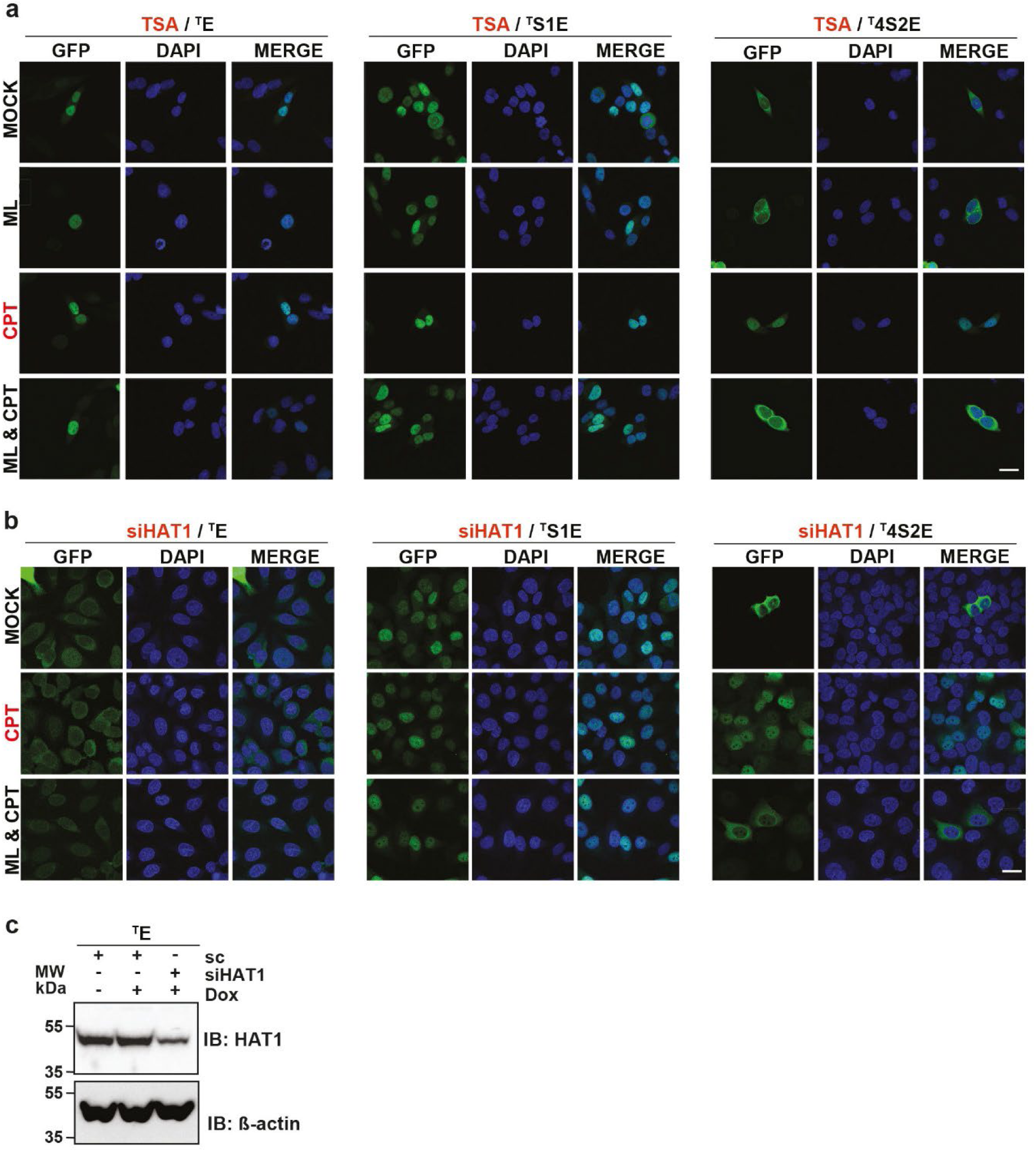
HAT1 and acetylated Histone H4 escort EME1 to DNA lesions. Multi-cell images and RNAi controls related to Fig. 4. **(a)** Multi-cell images associated to Fig. 4d. **(b)** Multi-cell images associated to Fig. 4e. **(c)** Immunoblot verifying RNAi mediated knock down of HAT1 for 48 hours in ^T^E expressing U2OS cells. Related to Fig. 4e. 32 hours post siRNA transfection, cell line was induced by 4 ng/ml Dox concentration. Cell lysates were resolved on SDS-PAGE and HAT1 expression (anti-HAT1) was monitored in comparison to anti-β-actin levels.

**Extended Data Fig. 5.**
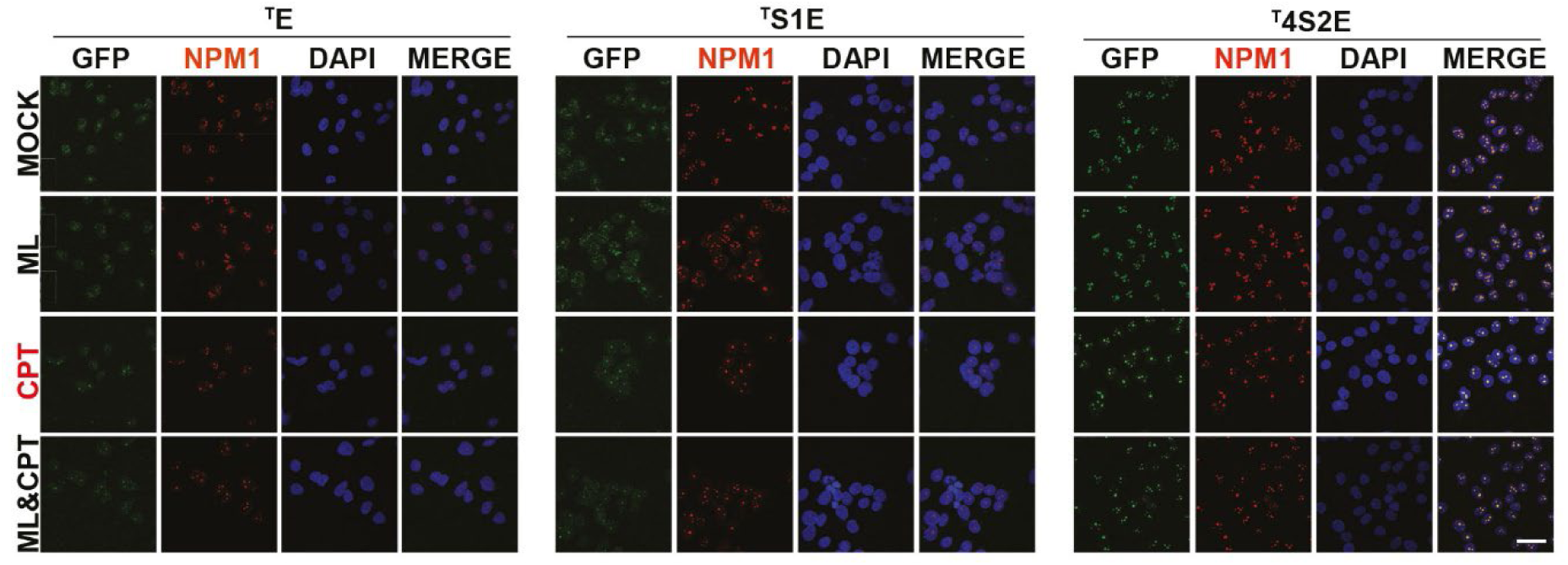
EME1 is sumoylated and retained at DNA lesions in nucleolar condensates. Related to Fig. 5. Multi-cell images related to Fig. 5c.

**Extended Data Fig. 6.**
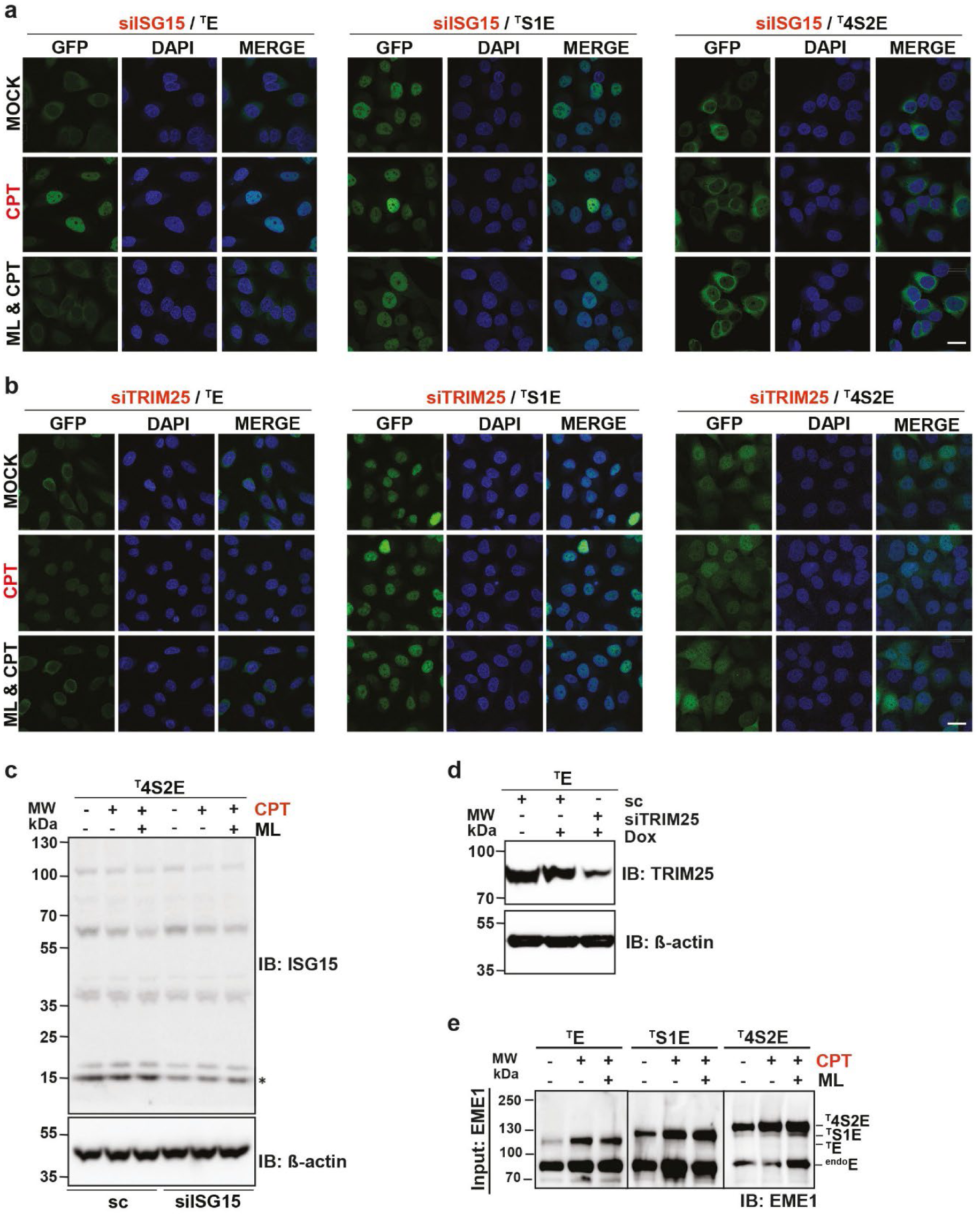
Removal of EME1 from DNA lesion via degradation and nuclear export & Endogenous EME1 is di- and poly-sumoylated upon CPT exposure. Multi-cell images and RNAi controls related to Fig. 6. **(a)** Multi-cell images related to Fig. 6c. **(b)** Multi-cell images related to Fig. 6d. **(c)** Immunoblot verifying RNAi mediated knock down of ISG15 for 48 hours in ^T^4S2E expressing U2OS cells related to Fig. 6c. 32 hours post siRNA transfection, ^T^4S2E expression for 16 hours and treated with DMSO, CPT and ML & CPT (as in Fig. 6c). Cell lysates were resolved on SDS-PAGE and ISG15 expression (anti-ISG15) was monitored in comparison to anti-β-actin levels. Star represents free ISG15. **(d)** Immunoblot verifying RNAi mediated knock down of TRIM25 for 48 hours in ^T^E expressing U2OS cells related to Fig. 6d. 32 hours post siRNA transfection, cell line was induced by 4 ng/ml Dox concentration. Cell lysates were resolved on SDS-PAGE and TRIM25 expression (anti-TRIM25) was monitored in comparison to anti-β-actin levels. **(e)** Input control to Fig. 6e.

**Extended Data Fig. 7.**
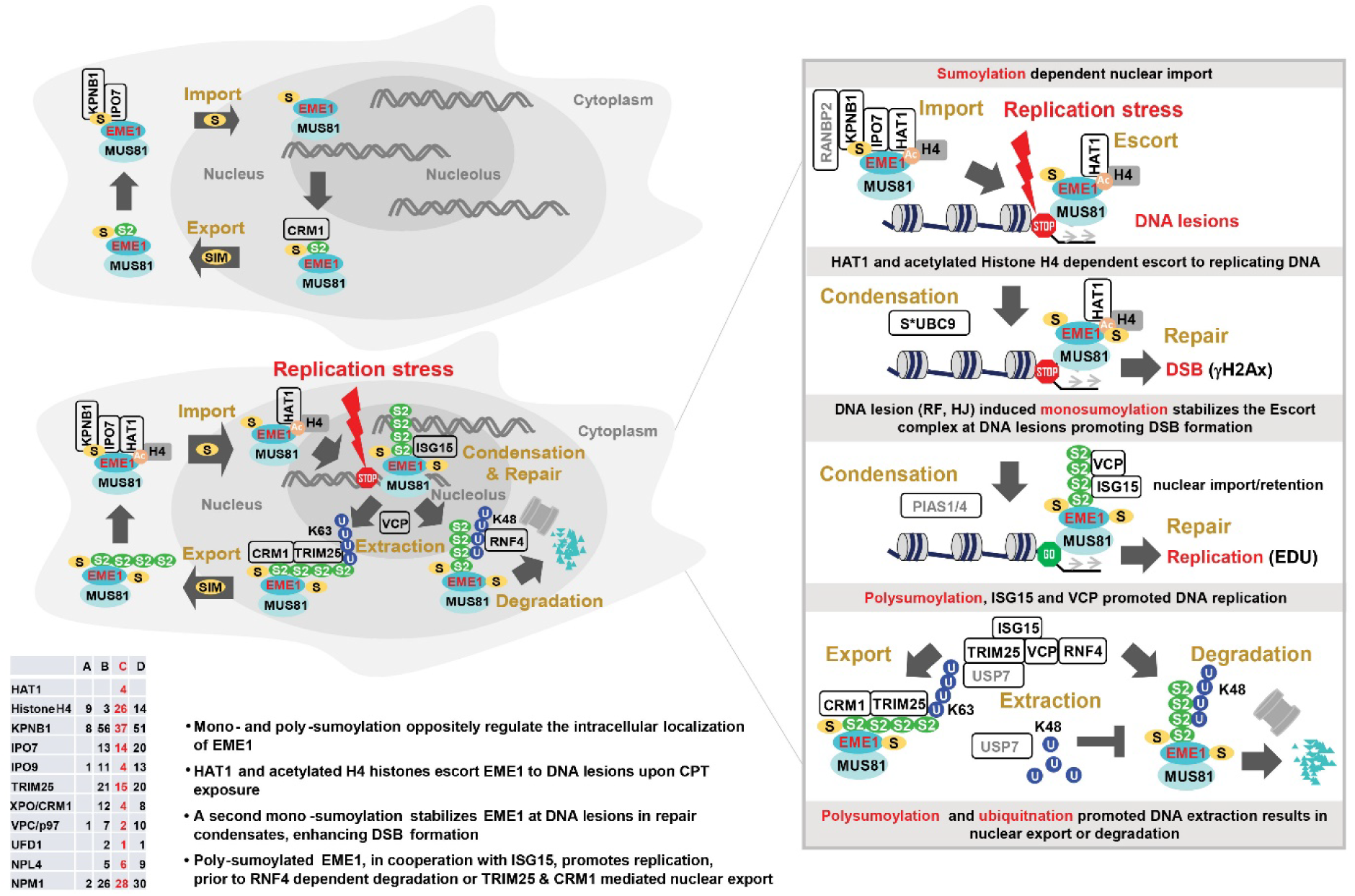
**(a) Model.** EME1 is a shuttling protein. In unstressed cells, nuclear export predominates nuclear import. EME1 nuclear import is dependent on KPNB1, IPO7 and mono-sumoylation, likely via RanBP2; nuclear export requires CRM1 and a functional SIM in EME1. Replication stress turns the transport ratio in favor of nuclear import, reinforced by HAT1 and acetylated H4 Histones that likely escort the EME1-MUS81 complex to sites of DNA replication. A second mono-sumoylation event, induced by the sumoylated form of UBC9 (S*UBC9), stabilizes EME1 in repair condensates in the presence of specific DNA structures (stalled replication forks or nicked Holliday junctions), which promotes the induction of resolving DSBs. Afterwards, EME1 is poly-sumoylated, likely by PIAS1/4, which facilitates DNA replication favored by the AAA ATPase VCP and ISG15. Then, poly-sumoylated EME1 gets extracted by the VCP-NPL4-UFD1 complex, a step that likely requires poly-sumoylation-dependent ubiquitination, either by RNF4 assembling K48-linked ubiquitin chains leading to EME1 proteasomal degradation or by TRIM25 conjugating K63-linked chains promoting nuclear export in cooperation with CRM1 and in dependence of the SIM of EME1. The SUMO chain conjugated to EME1 might interact with its SIM, leading to a closed conformation that is beneficial for nuclear export. Table presents a summary of peptide numbers for individual proteins supporting the proposed model under the indicated conditions obtained from EME1 IP-MS shown in Fig. 3a. IgG (A) and EME1 immunoprecipitations (IP) upon mock (B), CPT (C) and & ML & CPT (D) treatments. Key findings of this study are summarized in bullet points.

